# Modeling multiphage-bacteria kinetics to predict phage therapy potency and longevity

**DOI:** 10.1101/2022.11.11.516137

**Authors:** Zhiyuan Yu, Tiffany Luong, Selenne Banuelos, Andrew Sue, Mary Ann Horn, Hwayeon Ryu, Dwayne Roach, Rebecca Segal, Qimin Huang

## Abstract

*Pseudomonas aeruginosa* is a frequent cause of life-threatening opportunistic infections in the critically ill and immunocompromised. Its treatment is challenging due to the increasing prevalence of resistance to most conventional antibiotics. Although numerous alternative therapies are currently under investigation, bacteriophage (phage) cocktail therapy appears poised for long-term success. Here, we investigate potency and longevity of individual *Pseudomonas* phages in cocktail to determine viral co-factors that promote optimal treatment efficacy. We combined *in vitro* and *in silico* models to predict sixty-eight treatment permutations with three phages that adsorb symmetrically and asymmetrically when administered singly, double simultaneously, or double sequentially. We showed that simultaneously administering two asymmetrically binding phages with high cell lysis efficiencies improved cocktail potency. Use of a higher-potency cocktail, along with a reduction in the net probability of independent gene mutations was associated with prolonged bacterial suppression. Nevertheless, *in vitro* we almost always observed evolution of multiphage resistance. Simulations also predict that when combining phages with polar potencies, susceptible host cells are monopolized by the more efficiently replicating phage. Thus, further perpetuating the growth demise of the weaker phage in cocktail. Our mathematical model was used to explore and predict changes in phage and bacterial populations that were difficult to measure experimentally. This framework has many inferential and exploratory uses for clinical investigation such as identifying the most sensitive parameters for phage selection and exploring different treatment regimens. Collectively, our findings attempt to dissect the mechanisms of phage cocktails combating *P. aeruginosa* infections and highlight the viral co-factors necessary for treatment efficacy.

## 1. Introduction

Multidrug-resistance (MDR) is continuing to threaten global public health. The United States Center for Disease Control and Prevention (CDC) estimates that antibiotic resistance bacteria cause over 2.8 million illnesses each year, leading to tens of thousands of deaths as a result (CDC, 2019). *Pseudomonas aeruginosa* is a frequent and challenging nosocomial pathogen with consistently high rates of MDR in the US that ranges from 11.5% to 24.7% (1). MDR *P. aeruginosa* infections commonly occur in people in the hospital or with weakened immune systems. It is particularly dangerous for patients with chronic lung diseases like cystic fibrosis. MDR *P. aeruginosa* infection causes higher mortality, longer hospital length of stay, higher readmission rates, and US $20,000 excess cost per infection relative to those with non-MDR counterpart infection (2). While improving antibiotic stewardship has slowed the pace of MDR and there are new antibiotics in the development pipeline, newly marketed antibiotics will likely be quickly outpaced by the rapid emergence of resistance (3, 4).

As a century-old infection remedy, phage therapy is widely being redeveloped to treat MDR infections (5–7). Bacteriophages, or phages for short, are viruses that have evolved to infect and kill bacteria. As such, they pose limited adverse health effects in human cells and the body’s commensal microflora. Phages are primarily unaffected by common mechanisms of antibiotic resistance (5, 6). In fact, phage lytic pressure can restore antibiotic susceptibility in target bacteria via collateral sensitivity mechanisms (8). Preclinical data demonstrate that *Pseudomonas* phages exhibit high therapeutic potential when used alone or with standard of care antibiotics (9–13). In addition, phage therapy for *P. aeruginosa* has been employed in a limited number of individual patient expanded access investigational new drug (IND) applications (14–16). Nevertheless, there are disadvantages when administering phages as therapeutics. Most individual phages are highly strain specific, which provides clinical users a very narrow host range (5, 6). It is also common to observe the selection of phage resistance (10, 17–19). Expanding the host range and limiting resistance can be achieved by combining multiple phage types, each with different infection properties (11–13, 20). Because phages are also the most abundant form of “life” with incredibly large biodiversity, novel phages with unique infection properties can be easily isolated from the environment. Developing phage cocktails, however, has many challenges. It remains unclear how subtle differences in phage strain infection kinetics and viral replication properties can lead to differences in host lysis rates and evolution of phage resistance (21, 22). Also understudied is how different phage types react to one another in a mixed population. For instance, combining phage strains with different infection strategies may promote synergistic effects that enhance host cell lysis (i.e. potency) and/or prevent antiphage evolution. In contrast, there is the potential for phages to act antagonistically to one another by competing over bacterial host cells (23). Thus, phage strain selection and formulation of phage cocktails may significantly influence the efficacy of phage therapy.

Mathematical models can aid clinical investigations in a number of ways. Constructing a mathematical model requires critical consideration of biological mechanisms underlying the experimental work. Model development can reinforce the process of analyzing findings, reveal inconsistencies observed in the lab, or highlight gaps in results undetectable by experimental instruments. For example, designing a model for phage-bacteria studies must reflect that phages do not conform to linear kinetics. Because phages require bacterial hosts for replication, their growth is completely dependent on the host population. Recent studies combining *in vivo* data and mathematical modeling have uncovered key parameters that determine phage-immune synergy during therapy (10, 24). Other theoretical studies have investigated phage-antibiotic-immune synergy and phage-phage synergy to provide insight on developing combination therapies to treat MDR bacterial infections (25, 26). Despite understanding the synergistic effects between the phages and other treatment factors, basic understanding of the phage-bacteria kinetics remains incomplete. For kinetics between bacteria and one phage strain, Payne and Jansen developed a simple mathematical model to explore the initial dynamics of phage infection (27). Following Payne and Jansen’s work, Cairns et al. extended the work by including resistance to phage (28). However, the kinetics between bacteria and two phage strains has not been directly addressed and how phage resistance develops during multiphage treatment remains unclear. In addition, phage selection, dose timing, and collateral resistance during single and multiphage treatments are poorly understood.

In this study, we developed novel multiphage-bacteria mathematical models to explore the treatment efficacies of single, double simultaneous, and double sequential administration strategies with three *Pseudomonas* phages. We developed *in vitro* and/or mathematical models of time-kill kinetics of single and pairwise phages to show 68 permutations of single, double simultaneous, or double sequential treatment strategies. We then compared treatment strategies to delineate the influence of individual phage strains during these treatment strategies to suppress populations of *P. aeruginosa*, as well as limit the evolution of phage resistance. Moreover, our double phage mathematical models were extended to explore alternative application scenarios, such as manipulating dose and sequential timing. Ultimately, we found that phage potency and receptor asymmetry were the major co-factors to effective cocktail treatment and that double simultaneous treatments impose the highest fitness cost. Together, this work provides insights to improve phage cocktail therapy formulation and regimens.

## 2. Methodology

A variety of microbiological assays exist to test phage infection and bacterial responses to infection ranging from plaque formation assays to dynamic microfluidic devices (29–32). Our experiments use microplate time-kill assays, a gold standard to study the activity of antimicrobial agents *in vitro*, to demonstrate phage lytic and replicative properties and evolution of host bacteria to phage infection. The method uses a high-throughput 96 microwell format to measure bacterial growth under different phage exposures and conditions. In this work, we explore phage infection dynamics for the Gram-negative *P. aeruginosa* strain PAO1. We first performed a series of time-kill assays using 3 virulent (i.e. strictly lytic) phage strains, LUZ19, PYO2, and E215, which rapidly and efficiently lyse strain PAO1 *in vitro*. Characterization of single phage treatments as well as double phage cocktail treatments were compared to demonstrate phage-bacteria outcomes under the different treatment regimes. Using changes in bacterial density (measured by optical density (OD_600_)) we observed trends in bacterial killing and bacterial regrowth in the form of evolved phage resistance against one or more phage strains.

Next, we used the time-kill kinetics results and other known phage replication properties to develop and parameterize a mathematical framework for investigating the complexities of phage-bacteria interactions that were not resolved during *in vitro* experimentation. For example, the growth dynamics of individual phages when they were combined as a double phage cocktail treatment. We developed a single-phage ODE model and extended it to two versions of the double phage ODE model.

### 2.1 Experiment design and data description

#### 2.1.1 Bacteria and phage strains

*P. aeruginosa* strain PAO1 was grown aerobically in Luria broth (LB) Lennox at 37^◦^C with shaking (120rpm). Phages were propagated as previously described (33). Briefly, purified phages were added to mid-log growing bacteria at the multiplicity of infection (MOI) 0.1 and incubated overnight at 37^◦^C with shaking. Lysates were centrifuged at 8000×g to remove bacterial debris and the collected supernatant was 0.22 μm dead-end filter sterilized. Phage preparations were quantified by serial dilution spot titration and stored at 4^◦^C (33).

Virulent phages LUZ19, PYO2, and E215 are double-stranded DNA tailed viruses that were all originally isolated on the *P. aeruginosa* strain PAO1 (9, 34). Phage LUZ19 is a podovirus in the genus *Phikmvvirus* and has a 43,548-bp genome (34). LUZ19 infection consists of binding to the pili of the host cell with an adsorption rate of approx. 2.48 × 10^−10^ mL^−1^ min^−1^ (Supplementary Table 1) and a replication latent period of 17min, which produces a burst size of ~120 virions (35). PYO2 is a myovirus in the genus *Litunavirus* and has a genome size of 72,697-bp (9). PYO2 infection consists of binding to the outer membrane lipopolysaccharide (LPS) with an adsorption rate of approx. 1.8 × 10^−10^ mL^−1^ min^−1^ (Supplementary Table 1, Supplementary Figure 1) and a replication latent period of 20min, which produces a burst of ~200 virions. E215 is also a myovirus in the genus *Pbunavirus* and has a genome size of 66,789-bp (9). Of the three, E215 binds to its host the fastest at an adsorption rate of 1.4 × 10^−9^ mL^−1^ min^−1^ using LPS as the cell surface receptor. However, E215 exhibits the longest latent period of 30min and largest burst size exceeding >200 virions (Supplementary Table 1, Supplementary Figure 1).

#### 2.1.2 Phage titering

The spot titering method was used for phage quantification prior to each experiment (33). Briefly, *P. aeruginosa* PAO1 was grown to OD_600_ 0.2 and plated (lawned) onto dry LB agar plates and sterilely dried. Phage stocks were tenfold serially diluted using a microplate (10^−1^ to 10^−8^) and a multichannel pipette was used to spot phages onto the dried bacterial lawn. After sterilely drying phage spots, plates were statically incubated at 37°C overnight. Visible plaques were enumerated as plaque-forming units (PFU).

#### 2.1.3 Adsorption and one step

During phage replication, phages bind to the host (adsorption), hijack their cellular machinery to create phage proteins and assemble infectious phages (eclipse period), then are released through lysis of the bacterial host (latent period). The first step of phage-bacteria dynamics is the attachment of phages onto the susceptible bacteria host. The speed at which a phage binds is the adsorption rate, which was determined by measuring the decrease in free-phage particles in a culture of bacteria (36). Briefly, 1mL of a freshly diluted phage solution (1×10^7^ PFU/mL) was mixed into 9mL of 37°C incubated LB containing exponentially growing *P. aeruginosa* PAO1 at a concentration of ~5×10^7^ colony-forming units per mL (CFU/mL). As a result, the initial phage concentration during the experiment was 1×10^6^ PFU/mL to achieve a MOI of ~0.02. The mixture was incubated at 37°C with shaking (120rpm) and 50µL samples were taken every minute and vortexed with chloroform to forcibly lyse bacterial cells, leaving unbound free phage in the solution. All samples were centrifuged at 12,000×g for 3min to remove bacterial debris and to separate aqueous chloroform from samples. Phage titering was performed as previously described to quantify free phages at each timepoint.

To determine the number of phage particles produced during each replication cycle, called burst size, we performed the one-step growth curve protocol described previously in (37). Briefly, 1mL of a freshly prepared phage solution (5×10^7^ PFU/mL) was mixed into 9mL of 37°C incubated LB containing exponentially growing PAO1 (5.5×10^7^ CFU/mL). As a result, the phage concentration during the experiment is 5×10^6^ PFU/mL to achieve a MOI of 0.1. At 2min intervals, two 100µL samples were taken, with the first sample used to determine the total number of infected bacteria and the second treated with 50µL of chloroform to determine the number of phage particles. The chloroform-treated samples will induce premature lysis of the bacterial cells and measure the increase of free phage over time. Both sets of samples were spot titered as previously described. Titer comparison between chloroform treated and untreated samples was used to determine the eclipse and latent periods as well as the burst size. Adsorption and one-step growth analyses were performed in triplicate for each phage strain.

#### 2.1.4 Time-kill kinetics

Time-kill kinetics were used to examine the dynamics of phage mediated bacterial killing over time using a 96-well microplate format. As a measure of phage killing, the OD_600_ of bacterial cultures was measured over time and *P. aeruginosa* PAO1 growth under aerobic plate reader conditions for 28h was used to describe ancestral bacterial growth (no phage control). *In vitro* quantification of phage killing using a CLARIOstar microplate reader (BMG Labtech, Ortenberg, Germany) had a limit of detection (LoD) of OD_600_ 0.2. For each experiment, PAO1 inoculum was freshly prepared by growing a single CFU overnight in LB broth. Overnight cultures were diluted into fresh LB broth to OD_600_ 0.2 and grown for 2h to obtain bacteria in the early logarithmic phase of growth. Microplate wells were filled with 170μL of LB broth, inoculated with ~ 2 × 10^6^ CFU, and dosed with phages (MOI 1), before being sealed with a BreathEasy^TM^ membrane (Diversified Biotech, Dedham, MA, USA) to allow gas exchange. A microplate reader incubated the microplates at 37^◦^C under shaking conditions (120 rpm), while measuring OD_600_ at 6min intervals between shaking steps. Following incubation for 28h, individual microplate wells were streaked onto LB agar to isolate bacteriophage insensitive mutants (BIMs). Individual CFUs were subcultured, tested for phage resistance (see below), and archived with 25% glycerol at −80°C.

#### 2.1.5 Phage resistance assay

To assess phage resistance, BIMs were grown in LB broth to OD_600_ 0.2 and the spot titering technique was then performed as previously described, using BIMs for the bacterial lawn. The absence of phage plaques after 24h incubation at 37°C indicated phage resistance to the phage.

#### 2.1.6 DNA extraction and genome sequencing

In order to determine phage strain specific mutational differences in BIMs, DNA sequencing was performed on phage resistant CFUs isolated from single or cocktail treatment. Bacterial pellets were prepared by centrifuging bacterial liquid cultures at 8000×g and genomic DNA was extracted by NucleoSpin^TM^ Microbial DNA kit (Macherey-Nagel, Duren, Germany). Briefly, bacterial cells were physically disrupted by bead-beating and DNA was bound to silica spin-columns. Following washing (to remove salts and unwanted proteins), DNA was eluted into water, quantified, and sent to SeqCenter (Pittsburgh, PA, USA). Sequencing libraries were prepared by SeqCenter using the Illumina Nextera Kit and then sequenced on the Illumina NextSeq 550 platform (Illumina, San Diego, CA, USA) with paired-end 150bp nucleotide reads.

#### 2.1.7 Bioinformatic analysis

Because short-read sequencing was performed, bioinformatic tools were required to assemble bacterial genomes. Raw reads were trimmed by fastp v.0.20.1 (38) (quality control of sequencing reads and removal of low quality reads) and *de novo* assembled using SPAdes v.3.15.1 (39) to obtain larger contigs (long consensus region of DNA from short reads) for further analysis. Reference guided assembly was conducted using MeDuSa v1.6 using the *P. aeruginosa* PAO1 NCBI reference genome (NC002516.2) (40). Annotation of open reading frames (ORFs) to determine gene functions was conducted using Rapid Annotations of Subsystems Technology (RAST) v2.0 (41). This full assembly and annotation process was performed for ancestral PAO1 grown under microplate conditions for 28h and served as the reference for mutational analyses of phage resistant BIMs. For BIM analyses, fastp v.0.20.1 was used to generate quality controlled (clean) short reads. Variant calling was conducted using the breseq v.0.35.5 pipeline (42) to compare clean BIM short reads to ancestral PAO1. Default settings were used for all software analyses.

#### 2.1.8 Resistance fitness assay

Phage resistant populations of *P. aeruginosa* PAO1 were revived from the glycerol stocks, archived as previously described from the time-kill assay. A single CFU of each population was inoculated in 5mL LB test tubes and incubated at 37°C until the early logarithmic growth phase (OD_600_ 0.2). Cultures were centrifuged at 8,000×g for 5mins and resuspended in water for a concentration of 2×10^7^ CFU/mL. 96-well microtiter plates were filled with 100µL of fresh 2X LB and inoculated with 100µL of the bacteria-water suspension for a total of ~2×10^6^ cells per microwell. 96-well plates were sealed using BreathEasy^TM^ membrane and incubated in a microplate reader at 37^◦^C for 18h. OD_600_ measurements were taken at 6min intervals after a 120rpm orbital shaking step. Resistance fitness growth curves were repeated 3 times with individual CFUs from the archived stock.

### 2.2 Bacterial Growth Mathematical Model

We organized the *in vitro* data based on the treatment regime. Each dataset allows us to estimate key parameters in our mathematical models.

*D1.*Control: *P. aeruginosa* PAO1 growth without the addition of phage.

*D2.*Total bacterial density from single phage treatment: *P. aeruginosa* PAO1 growth after single phage treatment with either phage LUZ19, PYO2, or E215.

*D3.*Resistant bacterial density from single phage treatment: *P. aeruginosa* PAO1 BIM growth for each experiment described in D2.

*D4.*Total bacterial density from simultaneous double phage cocktail treatment: *P. aeruginosa* PAO1 growth after simultaneous double-phage cocktail treatment with a pair of phages from LUZ19, PYO2, and E215.

*D5.*Resistant bacterial density from simultaneous double-phage treatment: *P. aeruginosa* PAO1 BIM growth for each experiment described in D4.

#### 2.2.1 Single-phage Mathematical Model

For single strain phage therapy, Payne and Jansen developed a simple mathematical model to explore the initial dynamics of phage infection without accounting for phage induced bacterial changes (27). Following Payne and Jansen’s work, Cairns et al. extended the work by including phage resistance as a model parameter (43). Inspired by their work, we have developed a nonlinear four-compartment ODE model that describes the density-dependent interactions between ancestral bacteria, single phage treatment, and the emergence of phage resistant bacteria in an *in vitro* setting. The model variables are: the density of ancestral bacteria (original strain) (*B*), the density of phage-resistant single-mutant bacteria (*B_R_*), the density of phage infected bacteria (*B_I_*), and the density of phage (*P*). Figure 1 is a schematic depicting this interaction.

**Figure 1:**
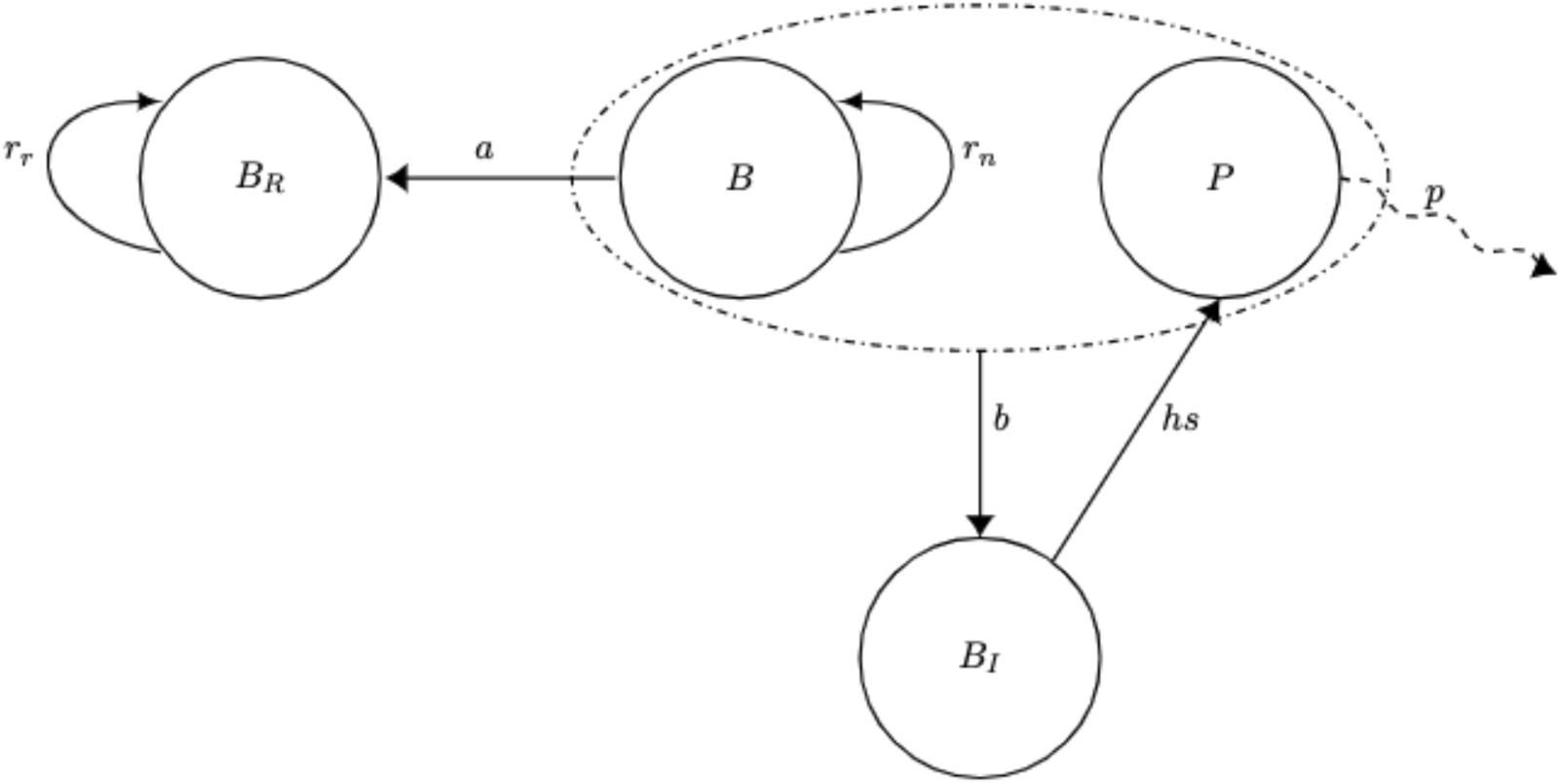
Schematic diagram for the in vitro single phage model. In the presence of phage (*P*), ancestral bacteria (*B*) either mutate into phage-resistant single-mutant bacteria (*B_R_*) or are infected by phage (*P*). Infected bacteria are subsequently moved to the infected class (*B_I_*). New phages are released when the infected bacteria cell is lysed.

Ancestral bacteria, *B*, exhibits logistic growth in the absence of phage with growth rate *r_n_*, carrying capacity *K*, and density dependence *B* + *B_R_*. As bacteria replicate with phages in their environment, mutations conveying phage resistance will emerge (17). We assume that ancestral bacteria mutate to resistant bacteria at a rate of *a*. Back mutation from *B_R_* to *B* was not included in the model because the probability of a reverse mutation is extremely low in the presence of high phage concentration selective pressures. Phage and bacteria are assumed to interact via mass action dynamics, thus we denote the binding rate of phage to bacteria as *b*. An infected bacteria cell can no longer replicate and will be lysed by the attached phage at a rate of *s*. The phage-resistant single-mutant bacteria, *B_R_*, grows logistically with growth rate *r_r_* and have no density loss due to phage infection nor mutation.

The growth in phage density is due to the release of phage progeny through lytic infection of *B_I_* at a burst size of *h*. We assume free phage particles are removed in the system by background decay at a rate *p*. In the *in vitro* setting this decay rate is quite small, so the effect of *p* is minimal. However, this term is kept in the model as it is significant for extended *in vivo* models and arises in the proliferation threshold.

The single phage model, model (1) is:

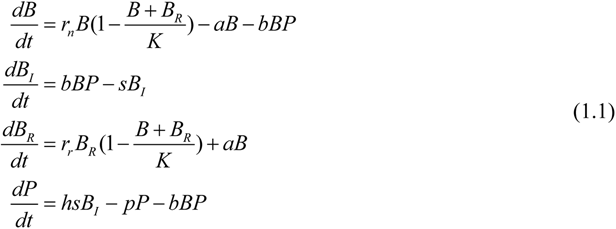

Since the rate of phage growth is dependent on the host population, Payne and Jansen introduced a proliferation threshold, which is the threshold density of bacteria that must be present in order for the phage density to increase (27). The reproduction number, the number of secondary infections arising from a single infected cell, must be larger than 1 for phage infection to propagate. Each infected bacterial cell’s lifespan is given by 1/*s* and produces 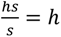 phage particles. In turn, each phage particle will cause 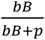 new infections. Thus, we need

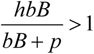

Hence, the proliferation threshold is given by

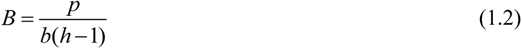

This means that the initial density of bacteria must be above the proliferation threshold in order for the phages to replicate effectively.

The inundation threshold is the threshold density of phage that will reduce the density of ancestral bacteria. Thus, we need 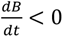.

We estimate

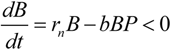

From here, we obtain the inundation threshold

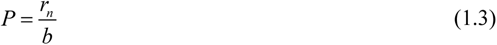

Although this mathematical model has been designed to describe an *in vitro* environment, the thresholds derived will be meaningful in determining timing and dose during phage therapy. Indeed, it will be important to confirm that the phage properties and patient bacterial loads meet the thresholds.

#### 2.2.2 Two-phage Mathematical Models

Using multiple phage strains during treatment is thought to be more effective than the application of a single phage type [12, 18]. Simultaneous treatment using two or more phage strains as a “cocktail” therapy may reduce the possibility of evolution of phage resistant bacterial mutants (higher selective pressure) and expand the host range of the treatment (5, 44). That is, the probability of two different resistance mutations occurring in a single host cell can be significantly lower than that of only acquiring a single mutation depending on the phage pairing (45). However, the kinetics of phage-phage-bacteria interactions are not well understood.

To the best of our knowledge, two-phage mathematical models have not been developed to explain the complex dynamics and demonstrate the effects of different treatment strategies on bacterial density and phage resistance emergence. For instance, is it more effective to design a treatment with different phage strains administered simultaneously or sequentially? Is there evidence of competition between different strains of bacteria?

To answer these questions, we propose two versions of the double-phage model. We deem this approach necessary because the types of resistant bacteria that emerge from cocktail treatment depends on the phage binding receptor(s) in the cocktail. In our experimental data PYO2, E215 and LUZ19 use the cell surface receptor of LPS, LPS, and pili, respectively. All the phages in a cocktail of PYO2 and E215 will bind to the same cell surface receptor and select for ‘collateral resistance’ in isolated BIMs. Conversely, a combination of PYO2 and LUZ19, or E215 and LUZ19 will select for BIMs with independent gene mutations. This will cause the bacterial population to contain three subpopulations with: (i) resistance to Phage 1, (ii) resistance to Phage 2, and (iii) resistance to both phage strains.

We calibrated the parameters for our two-phage models using data measured from simultaneous treatment dynamics (D4 and D5). We then extend our models to study sequential treatment dynamics by supplementing our data with parameters assumed from literature (11). In the following sections we present both models. For simplicity, we continue to assume that there is no reverse mutation as a mutation in the identical region of the previous point mutation is quite unlikely.

#### 2.2.3 Two-phage mathematical model without collateral resistance

In the two-phage model without collateral resistance, we extend the single-phage model (1.1) (Figure 1) to allow for two different phage strains, *P*_1_ and *P*_2_, and three strains of bacteria: phage strain 1-resistant single-mutant bacteria, *B_R_1__*, phage strain 2-resistant single-mutant bacteria, *B_R_2__*, and double-mutant bacteria, *B_R_12__*. In addition, moving forward we define the “double-mutant strain” as a bacterial strain with mutations in both the pili and LPS loci. This model is suitable to describe, for example, the dynamics occurring when phages with different phage resistance outcomes are combined. The interactions between each phage and phage specific resistant bacteria are similar to the single-phage model. Figure 2 illustrates the corresponding schematic diagram.

**Figure 2:**
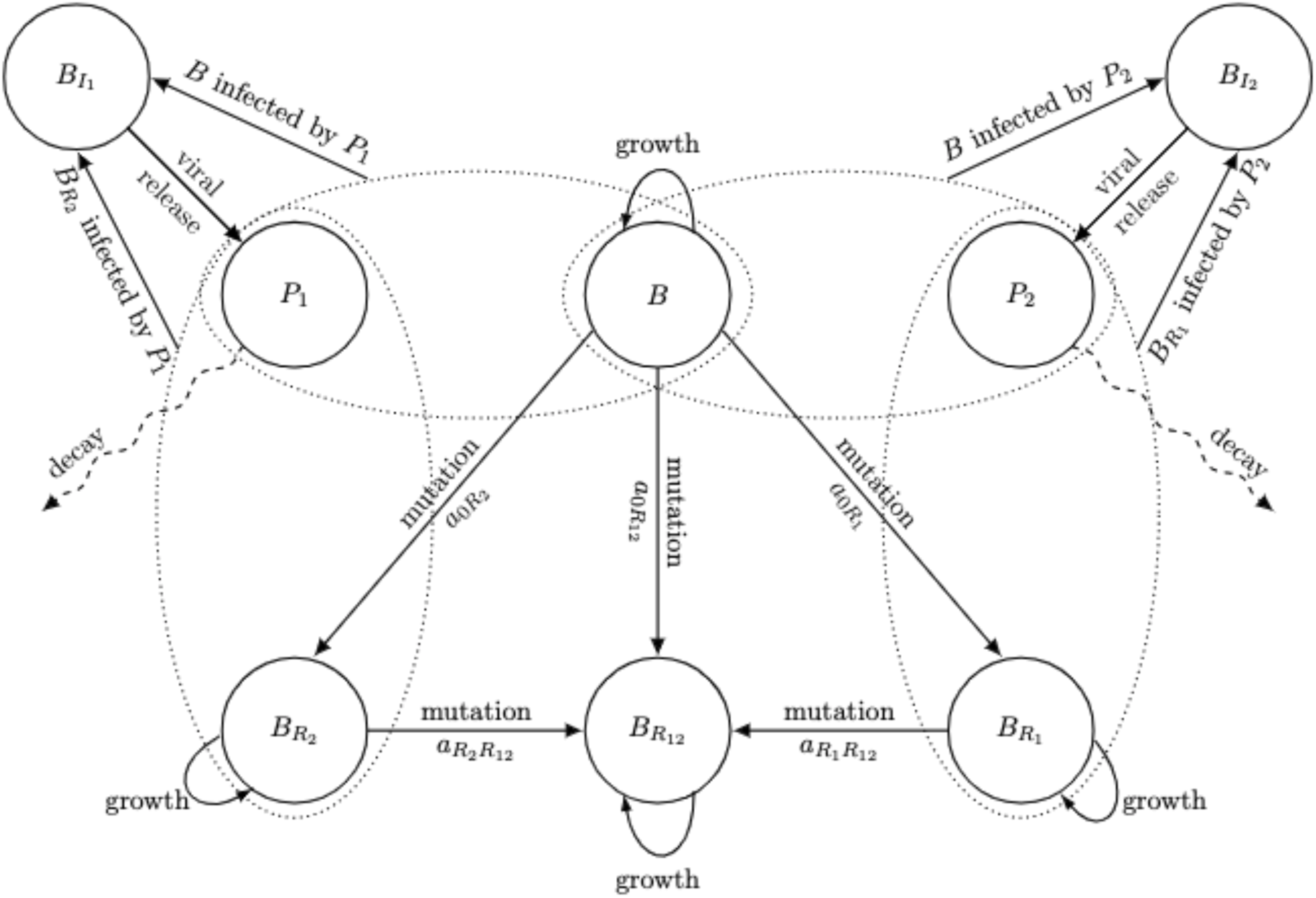
Schematic diagram for the in vitro two-phage model without collateral resistance. In the presence of phage (*P*_1_ or *P*_2_), ancestral bacteria (*B*) either mutate to resistant bacteria (*B_R_1__*, *B_R_2__*, or *B_R_12__*) or are adsorbed by phage (*P*). Adsorbed bacteria are subsequently moved to the infected class, (*B_Ii_*).

In the absence of phage, the ancestral bacteria, *B*, phage strain *j*-resistant bacteria (*j* = 1,*2*), and double phage resistant bacteria grow logistically with density dependence *B_T_* = *B* + *B_R_1__* + *B_R_2__* + *B_R_12__*, carrying capacity *K*, and growth rate *r_n_*, *r_j_*, and *r*_12_, respectively,.

In the presence of phage, the ancestral bacteria are either infected by phage strain *P_j_*, at rate *b*, acquire resistance to one of the phage strains at rate *a*_0_*_R_j__* or acquire a double-mutated gene with resistance to both phage strains at rate *a*_0_*_R_12__*. As explained above, this model is applied when one phage is pili-binding (LUZ19) and the other is LPS-binding (PYO2 or E215). Thus, if *a*_0_*_R_1__*: = *a_p_* then *a*_0_*_R_2__*: = *a_l_*. These parameter values are the same as in Table 1. Also, bacteria that has resistance to both strains of phages is denoted by *a*_0_*_R_12__*: = *a_lp_* in Table 2.

**Table 1:**
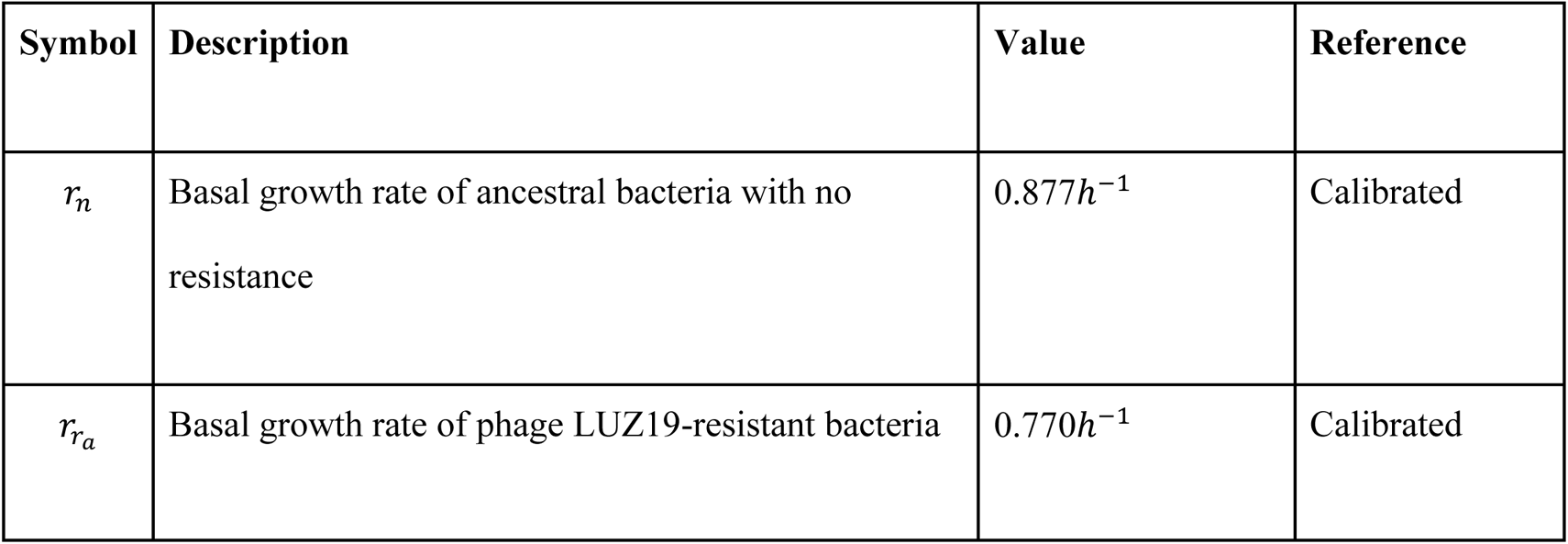
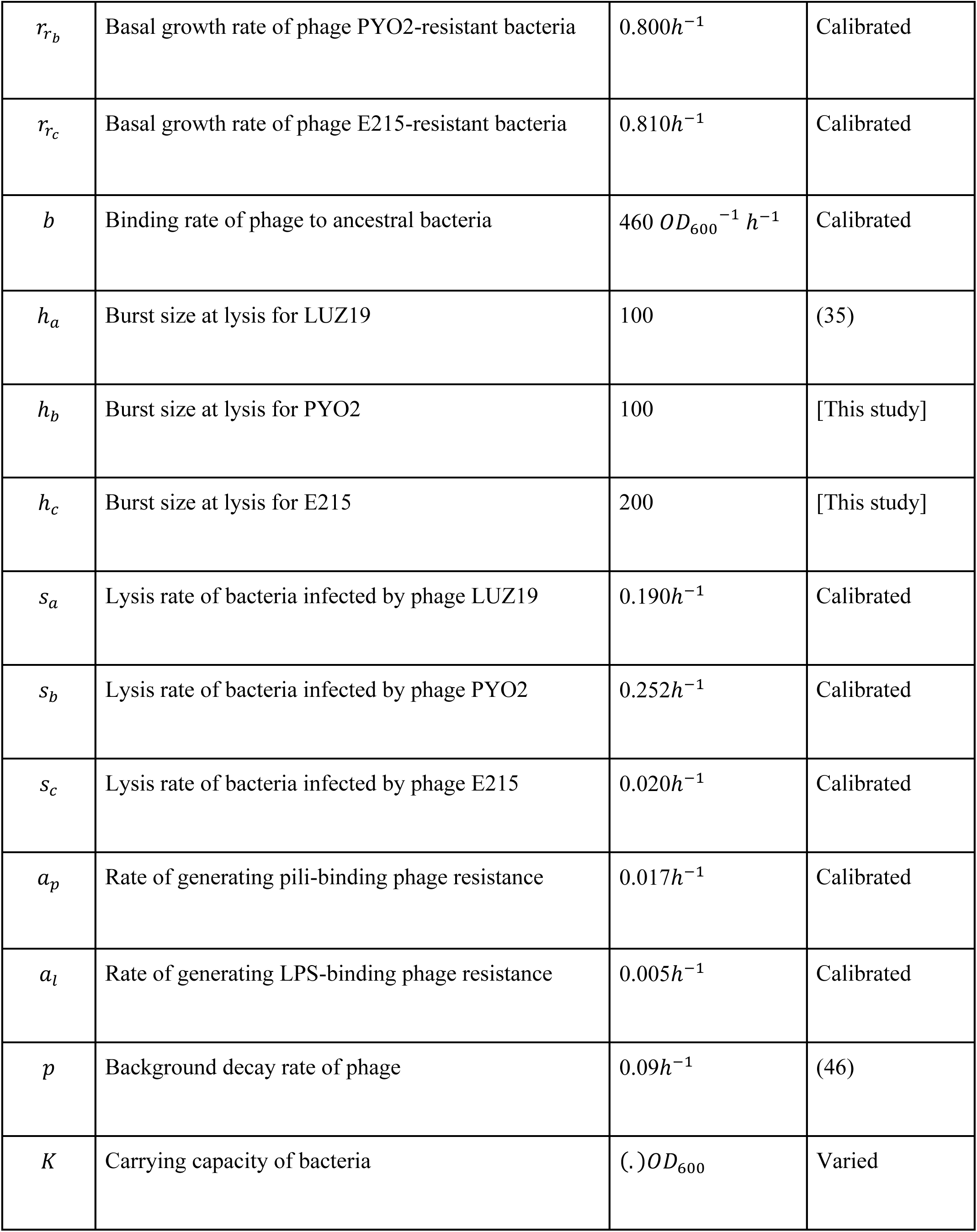
Parameters in single phage model

**Table 2:** Parameters new to the two-phage model

| Symbol | Description | Value | Reference |
| --- | --- | --- | --- |
| $a_{lp}$ | Rate of generating both pili-binding phage resistance and LPS-binding phage resistance simultaneously | $7.2 \times 10^{-6} h^{-1}$ | Calibrated |
| $r_{a+b}$ | Basal growth rate of bacteria with resistance to both phage LUZ19 and PYO2 in simultaneous simulation | $0.540 h^{-1}$ | Calibrated |
| $r_{a+c}$ | Basal growth rate of bacteria with resistance to both phage LUZ19 and E215 in simultaneous simulation | $0.618 h^{-1}$ | Calibrated |
| $r_{b+c}$ | Basal growth rate of bacteria with resistance to both phage PYO2 and E215 in simultaneous simulation | $0.800 h^{-1}$ | Assumed equal to $r_{r_b}$ in Table 1 |
| $r_{a \rightarrow b}$ | Basal growth rate of bacteria with resistance to both phage LUZ19 and PYO2 in phage (LUZ19→PYO2)- | $0.486 h^{-1}$ | (11) |
|  | sequential simulation |  |  |
| $r_{b \rightarrow a}$ | Basal growth rate of bacteria with resistance to both phage PYO2 and LUZ19 in phage (PYO2→LUZ19)-sequential simulation | $0.540h^{-1}$ | (11) |
| $r_{a \rightarrow c}$ | Basal growth rate of bacteria with resistance to both phage LUZ19 and E215 in phage(LUZ19→E215)-sequential simulation | $0.556h^{-1}$ | (11) |
| $r_{c \rightarrow a}$ | Basal growth rate of bacteria with resistance to both phage E215 and LUZ19 in phage(E215→LUZ19)-sequential simulation | $0.618h^{-1}$ | (11) |
| $r_{b \rightarrow c}$ | Basal growth rate of bacteria with resistance to both phage PYO2 and E215 in phage(PYO2→E215) - sequential simulation | $0.800h^{-1}$ | Assumed equal to $r_{r_b}$ in Table 1 |
| $r_{c \rightarrow b}$ | Basal growth rate of bacteria with resistance to both phage E215 and PYO2 in phage (E215→PYO2)-sequential simulation | $0.800h^{-1}$ | Assumed equal to $r_{r_b}$ in Table 1 |

Phage strain *j*-resistant single-mutant bacteria, *B_R_j__*, are infected by phage strain *P_j_* (*i* ≠ *j*), at rate *b*. The double-mutant bacteria, *B_R_12__*, are not infected by any phage. Phage strain *j*-resistant single-mutant bacteria mutate to double-mutant bacteria at rate *a_R_j_R_12__*. Given our possible scenarios, if *a*_0_*_R_1__*: = *a_l_*, then *a*_R1R2_ : = *a_p_*, for example.

Bacteria bound by phage strain *P_j_* move to the *B_Ij_* class. As in system (1.1) (See Figure 1), the infected bacteria no longer replicate and will lyse by phage strain *P_j_* at a rate of *s_j_*. The system of equations, model (2) is given by:

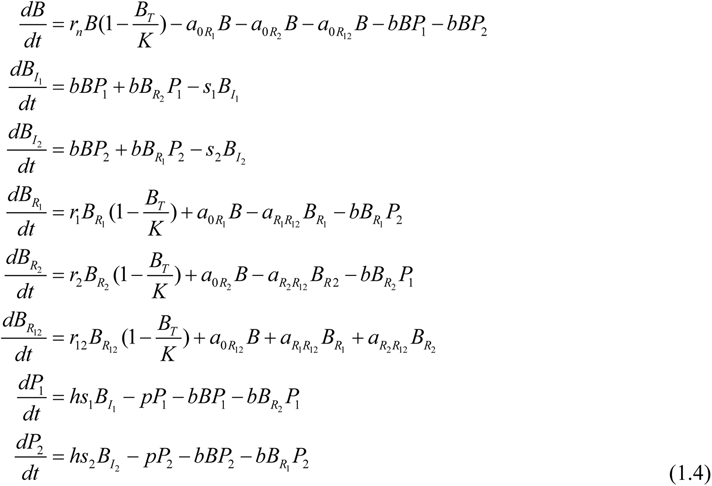

#### 2.2.4 Two-phage Model with collateral resistance

In the two-phage model with collateral resistance, only one bacterial strain emerges that is resistant to both phages, *B_R_*. This occurs when different phages target the same types of receptors on the bacteria cell wall. This model can describe the density-dependent interactions with experiments that applied phage PYO2 and phage E215, as both are LPS-binding phages. See Figure 3 for the schematic. The system of equations for this model is similar to system (2) without bacteria strains *B_R_1__* and *B_R_2__*. The system of equations, model (3) is defined as,

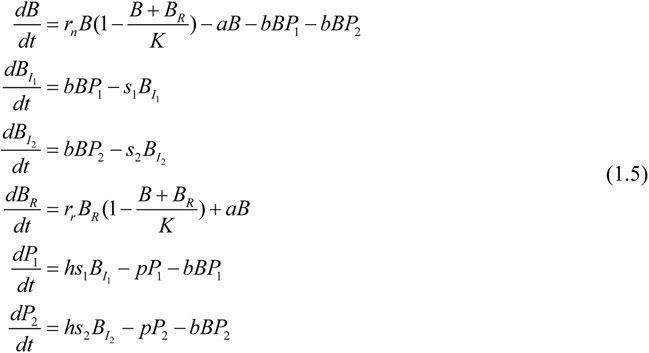

**Figure 3:**
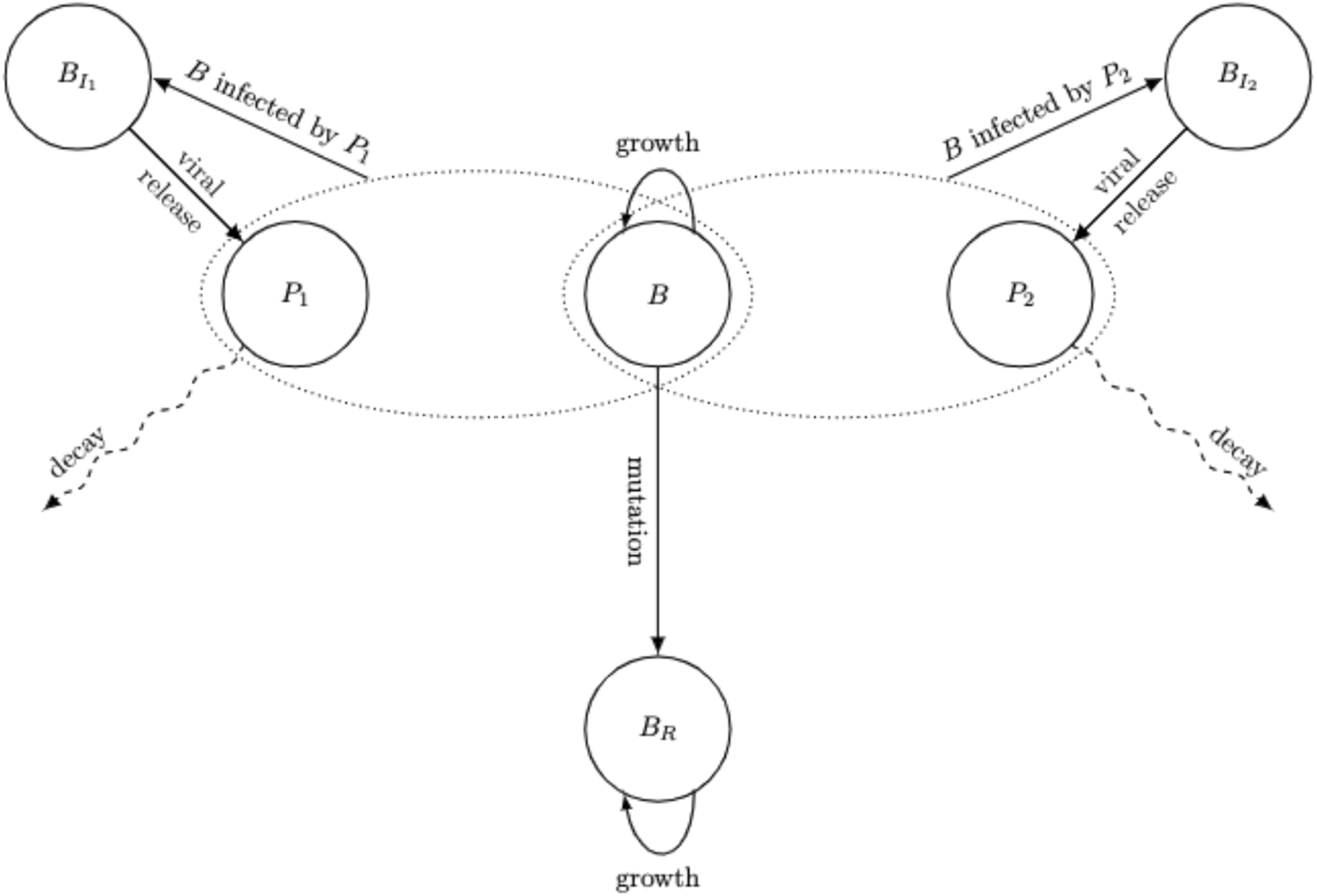
Schematic diagram for the in vitro two-phage model with collateral resistance. In the presence of phage (*P*_1_ or *P*_2_), ancestral bacteria (*B*) either mutate to bacteria with collateral resistance (*B_R_*) or are adsorbed by a phage strain. Phage-bound bacteria are subsequently moved to the phage strain infected class, (*B_I1_* or *B_I2_*).

## 3. RESULTS

### 3.1 Parameter estimation: Bacterial growth rate of *P. aeruginosa* PAO1

Individual cultures of *P. aeruginosa* were seeded into microculture wells and grown for 28h without phages to determine a baseline growth rate in the absence of phage infection. We used the average of 89 independent experimental growth curves to generate a logistic model to predict the growth of ancestral PAO1 (D1, Figure 4 blue curve). The theoretical limit of detection which correlated to OD_600_ 0.2 was calculated to contain 3.64 × 10^6^ CFU per well, corresponding to a concentration of 2.14 × 10^7^ CFU/mL.

**Figure 4:**
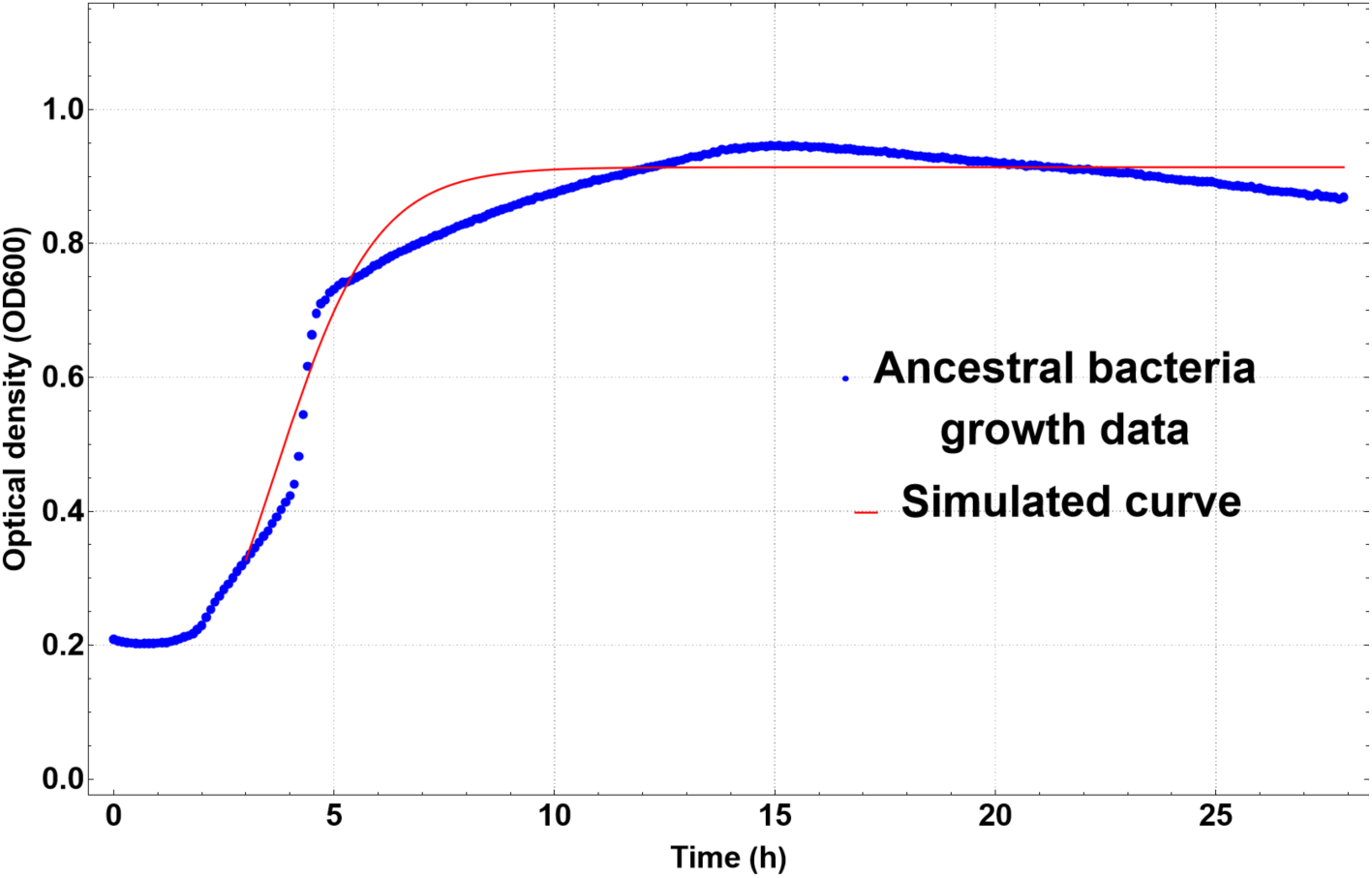
Ancestral bacterial growth of *P. aeruginosa* PAO1 without the addition of phage (blue curve) and logistic model fitting with Equation (1.6) (red).

We then calibrated the basal growth rate, *r_n_*, in the logistic growth equation for *B*, given by:

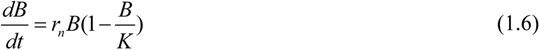

Figure 4 shows the result of this curve fit created using a least-squares fit with FindFit in Wolfram Mathematica.

### 3.2 Parameter estimation: Single phage model

*P. aeruginosa* PAO1 was treated in a time-kill assay with phages at 0h at a MOI of 1 and OD_600_ was measured (with shaking) for 28h. Treatment using individual phages LUZ19 and PYO2 successfully reduced the bacterial burden to the limit of detection within 0-4h (Figure 5A-B) while phage E215 did not reduce the bacterial burden below OD_600_ 0.2 in the same timeframe (Figure 5C). We therefore categorize LUZ19 and PYO2 as highly lytic phages with greater potency than E215. Exponential re-growth of the bacterial population occurred around 10-12h after phage addition, suggesting the presence of a phage resistant population. Following 28h of measurements, we isolated single colonies from each treatment condition. Characterization and sequencing revealed phage resistant *pil* (pili) mutants from LUZ19 treatment, and *wzy* (LPS) mutants from either PYO2 or E215 treatment. After characterization of the time-kill assay, we sought to mathematically model the observed dynamics and outcome. As shown in Figure 5, the predictions from the calibrated single-phage model were very close to the observed experimental data - *P. aeruginosa* PAO1 growth after single treatment with either phage LUZ19, PYO2, or E215. The parameters are listed in Table 1.

**Figure 5:**
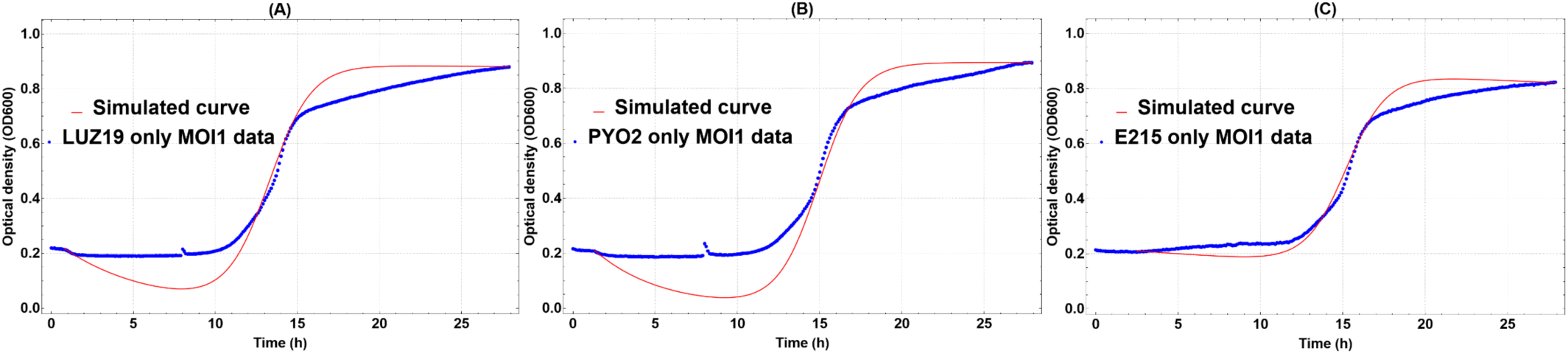
Growth curves of *P. aeruginosa* PAO1 treated with phages at MOI 1 (blue curve) at 0h with (A) LUZ19, (B) PYO2, and (C) E215. Model fittings (red) according to Equation (1.1). Note that OD600 0.2 is the detection limit of the plate reader.

Estimates for the parameter values of the binding, *b*, mutation, *a*, and lysis, *s*, rates for the system of equations of the single phage model (1.1) were calculated with FindFit and the data set D2. We make the following assumptions about the system for our calibrations:

1. Phages LUZ19, PYO2, and E215 are strictly virulent, meaning their reproduction is due to only a lytic lifestyle. We assume the binding of phage to bacteria will lead to infection, i.e. the binding of phage to bacteria is irreversible. Although there are differences in the binding mechanism depending on the phage characteristics, the binding term in our equation also includes rate of interaction between phage and bacteria, and due to the large number of cells, that process dominates the size of the rate term. Therefore, the binding rate of phage to bacteria is dominated by the physical contact of the bacteria and phage, and so different phage strains should effectively have the same binding rate *b*.
2. We also assume that different phage strains have different lysis rates, *s*, indicating their different potencies. The average lifespan of an infected cell initiated with the attachment of phage and ending with the bursting of the cell is given by 1/*s*. The phage specific lysis rates, *s_a_*, *s_b_*, and *s_c_* were estimated from the respective individual phage treatments. Consistent with the experimental latent period measurements for all three phages (Supplementary Table 1), the model-estimated phage specific lysis rates indicate that LUZ19 and PYO2 lyse the host cells faster than E215.
3. We were able to compute the bacteria growth rate from isolated growth experiments (Figure 6, Equation (1.6)) and estimate the mutation rate from time to appearance of resistance in the time-kill assay (Figure 5). There is no large difference in per gene mutation rate, denoted as *a* in Equation (1.1), on a molecular level across different strains of bacteria. However, the rate of generating a specific receptor mutation depends on the genetic basis of its phenotype. According to mutational analysis (of 20 sequenced phage treated isolates), there are 7 non-synonymous single nucleotide polymorphisms (nSNPs) that convey resistance to the pili-binding gene (Supplementary Table 2), but nSNPs occur in only one gene that encodes O-antigen used in LPS-binding (Supplementary Table 2). Thus, we assume that the rate of generating pili-binding phage (e.g. LUZ19) resistance, *a_p_*, is greater than the rate of generating LPS-binding phage (e.g. PYO2 and E215) resistance *a_l_*, i.e., *a_p_* > *a_l_*. Therefore, as shown in Table 1, the rate of generating pili-binding phage resistance (*a_p_*) is about three folds higher than the rate of generating LPS-binding phage resistance (*a_l_*), which explains why phage resistance arises faster in the phage LUZ19 only treatment compared to the other two single phage treatments (Figure 5).

**Figure 6:**
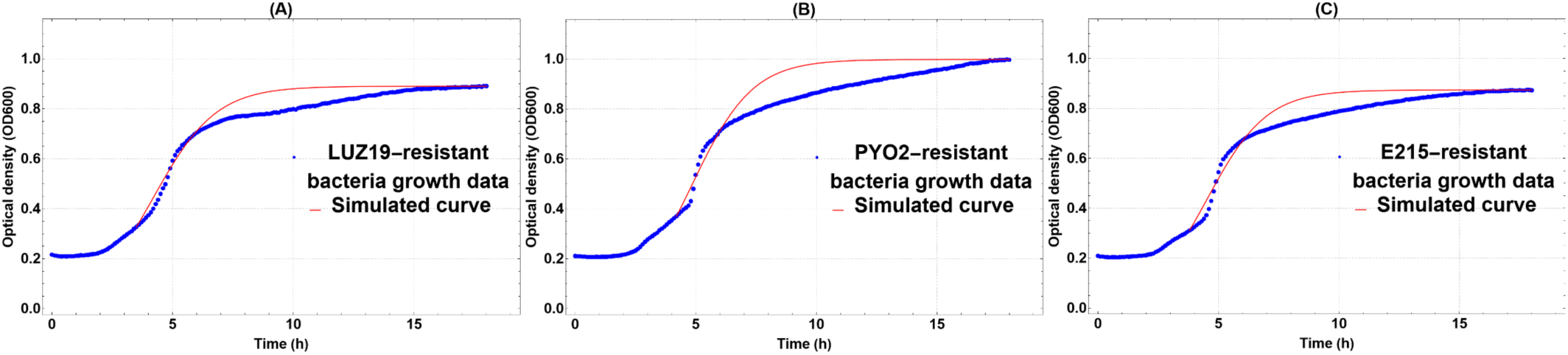
Growth curves (blue) of phage resistant PAO1 mutants to LUZ19 (A), PYO2 (B), and E215 (C) over 18h without phage treatment. Model fittings (red) according to Equation (1.6).

Finally, the growth rate, *r_r_*, for each single phage resistant strain that emerged from phage treatment is estimated by fitting Equation (1.6) to the respective data - Phage resistant *P. aeruginosa* growth data, isolated from each phage of the LUZ19, PYO2, and E215 treatments. Figure 6 shows the result of this curve fit created using a least-squares fit with FindFit in Wolfram Mathematica. However, the remaining parameters in the model either represent combination actions or are not measurable in isolation from experiments. We use the single phage model (equation (1.1)) along with the experimental data to estimate the remaining key parameter values such as mutation rate, binding rate, lysis rate (Table 1).

### 3.3 Model solution: Single phage model

For our *in vitro* experiments, we used OD_600_ as an approximation of phage-bacteria kinetics but only bacterial density can be measured spectrophotometrically. Thus, in order to convert our *in vitro* data into discrete CFUs for our mathematical model and to derive values for phage density, we calculated a bacterial standard curve to estimate CFU per/mL in the plate reader. We approximated that the OD_600_ is converted to CFU/mL by using *y* = 5*7*115528.3*7x* + 9*7*854*70*.84, where x is the OD_600_ value. Using these values and the parameters we calibrated for phage characteristics, we estimated bacterial CFU/mL and phage PFU/mL in our models.

For the single phage model (Figure 7), we model an equal ratio of ancestral bacteria and phages. Initially, the ancestral bacteria population decreases as the phage density is above the inundation threshold (Equation (1.3)). Meanwhile, the phage population surges as the ancestral bacteria density initially remains above the proliferation threshold (Equation (1.2)). Eventually, the resistant bacterial population dominates. The system behaves differently when different phage strains are applied. In early hours, potent phage strains, LUZ19 and PYO2, proliferate and reduce the total bacteria density much faster than E215. In later hours, the model predicts that the densities of the potent phage strains shrink much faster than the density of the less potent phage strain.

**Figure 7:**
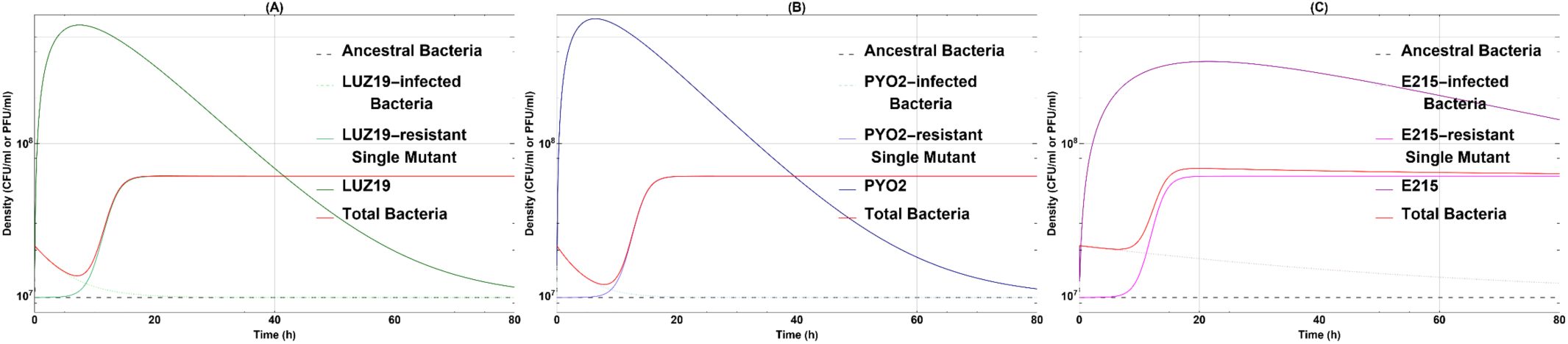
Single phage model numerical simulations: (A) LUZ19 only treatment, (B) PYO2 only treatment, and (C) E215 only treatment. For all the simulations, *B*(0) = 0.2, *B_I_* (0) = 0, *B_R_*(0) = 0, *P*(0) = 0.2, *b* = 460, *r_n_* = 0.877. For each simulation, phage specific parameter values can be found in Table 1.

### 3.4 Sensitivity analysis: Single phage model

We estimate the parameter values for our model from experimental data, but not all could be experimentally measured, so we studied the local sensitivity of the parameters to determine possible areas of concern. For example, the mutation rate is difficult to determine because bacterial density less than OD_600_ 0.2 is below the plate reader’s limit of detection. Our binding rate is a lumped parameter that includes rate of contact and adsorption time. To understand the relative effect changes in these parameter values have on the model transients, we perform local sensitivity analysis on each of the model states with the three phage strains. The method for this analysis is described in Appendix A and the results are shown below (Figure 8). The bacteria growth rate of the resistant bacteria, *r_r_*, and the lysis rate, *s*, are the most sensitive parameters locally. We further explore this by detailing the dependence of the model outcome on the phage specific lysis property.

**Figure 8:**
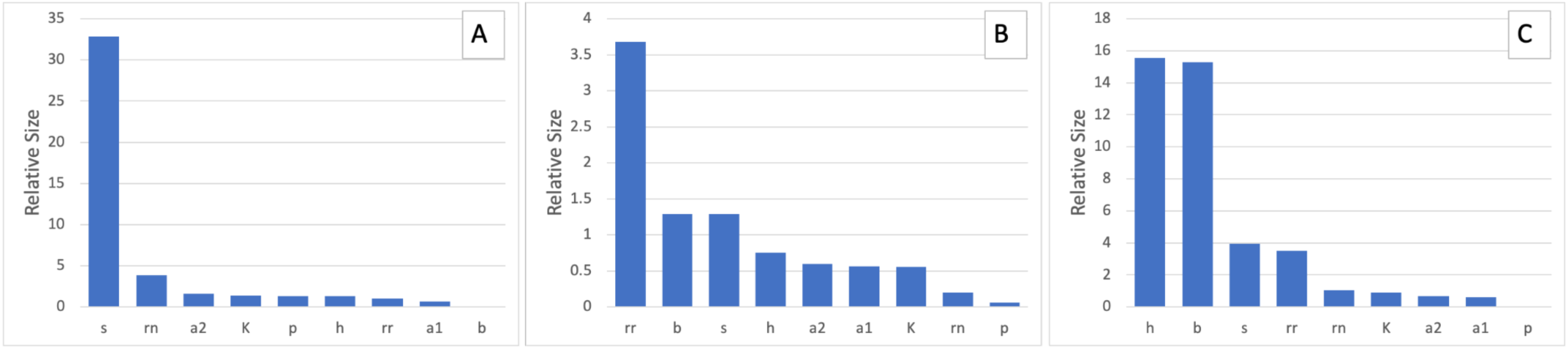
Sensitivity analysis for the single phage model of each phage parameter settings. Panel A has the relative sensitivities for phage LUZ19, panel B has the relative sensitivities for phage PYO2, and panel C has the relative sensitivities for phage E215.

By this measure, the only parameter deemed sensitive for all three phage strains is *s*, as shown in Figure 8. It is interesting to note that phage PYO2 and phage E215 also showed high sensitivity to *h*, *r_r_*, and *b*. Phage LUZ19 showed sensitivity to *r_n_*. Again, the mechanism of infection for phage LUZ19 differs from that of phage PYO2 and E215 and that difference is evident in the system activity.

### 3.5 Exploration of phage characteristics: single phage model

Using the parameters and observations we derived from *in vitro* experiments and *in silico* models above, we use the single phage model to explore hypothetical situations. There are millions of phages in existence, each with different properties. It is impossible to experimentally test them for efficacy against individual clinical pathogens. However, we can identify characteristics that would suggest that a phage is a good therapeutic candidate.

Since we used our model to define burst size and lysis rate as phage strain characteristics, we now explore the combination that may be most effective in delaying the emergence of phage resistant bacteria. For each parameter combination, we simulate the single phage model and find the time when the resistant bacteria density first exceeds OD_600_ 0.2 (the plate reader’s LoD). As shown in Figure 9, the phage with larger burst size and higher lysis rate tends to delay the occurrence of resistance more effectively. In addition, when the lysis rate is low, the time of occurrence of resistance is more sensitive to the change in lysis rate relative to the burst size. When the lysis rate is large enough, the time of occurrence of resistance becomes more sensitive to the change in burst size relative to the lysis rate. The three phages used in this study are indicated in Figure 9. PYO2 and LUZ19 are more effective in delaying the occurrence of phage resistance due to their high potency, or lysis rate. However, despite E215 having a burst size twice that of PYO2 and LUZ19, its low lysis rate constrains its ability to delay the occurrence of resistance.

**Figure 9:**
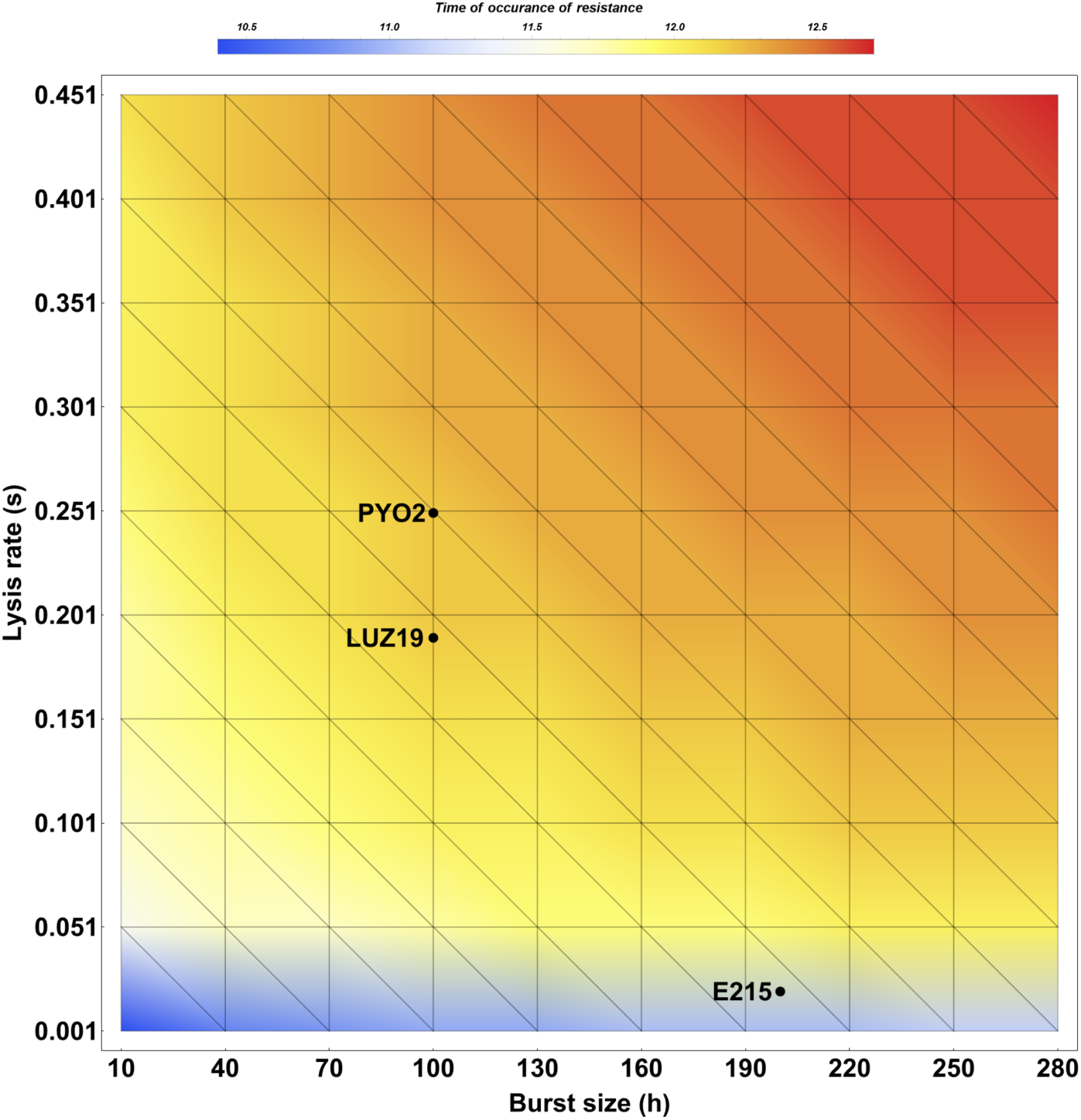
Different combinations of phage lysis rate (*s*) and burst size (*h*) and their corresponding time for the resistant bacteria to become detectable. The marked points denote our experimental phage strains PYO2, LUZ19, and E215.

### 3.6 Parameter estimation: two-phage model

The addition of multiple phage types against a single bacterial strain is desirable to exert sufficient killing pressure. Since increasing single phage MOI does not limit or prevent phage resistance, we examine the efficacy of three different two-phage cocktail treatments against *P. aeruginosa*. The same starting inoculum of strain PAO1 was used and a two-phage cocktail (MOI 0.5 per phage, total MOI 1) was added at 0h. Following the 28h incubation, we once again isolated and characterized CFUs from each cocktail treatment. Cocktail treatment using LUZ19 and PYO2 reduces the bacterial population to the lowest density in this study (Figure 10). Comparatively, cocktail treatment LUZ19 and E215 appears equally potent but resistance appears to emerge ~25h after treatment (Figure 10). Resistance to cocktail treatment PYO2 and E215 emerged the earliest ~12.5h, similar to the single phage treatment using either PYO2 or E215 (Figure 5). We found mutations against pili or LPS corresponding to the phages used (Supplementary Table 2). Both *pil* (pili) and *wzy* (LPS) mutations were identified in double-phage resistant mutants from the receptor-mixed cocktails while single SNP *wzy* mutants were sequenced from cocktail PYO2 and E215 treatment.

**Figure 10:**
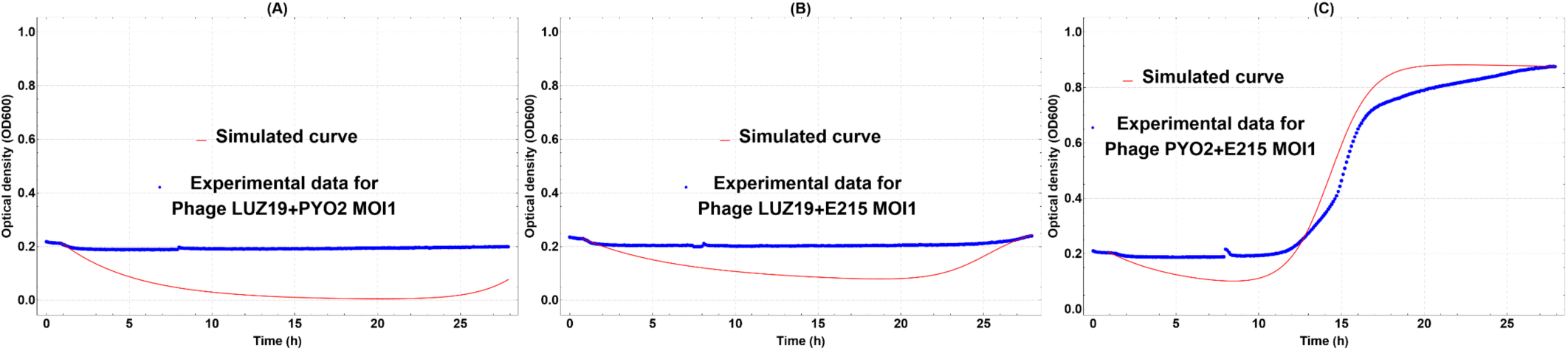
Growth curve (blue) of *P. aeruginosa* treated with cocktails (A) LUZ19+PYO2, (B) LUZ19+E215, or (C) PYO2+E215 over 28 h. Model fittings (red) to the experimental data. Note that OD600 0.2 is the detection limit of the instrument measuring the bacterial density.

Our two-phage treatment models (5) and (6) are used, in part, to simulate the interactions between phage and bacteria in an *in vitro* setting in order to study the effectiveness of different simultaneous treatments. Mathematically, the different treatment options require different parameter values in the growth rates of the double resistant bacteria that emerge. Experimental data D4 and D5 - *P. aeruginosa* PAO1 growth and phage resistant growth after simultaneous two-phage cocktail treatment with a pair from LUZ19, PYO2, and E215 - are utilized to calibrate the rates for these simulations.

As explained in Section 2.2 model (5) is utilized in simulations when phage LUZ19 is applied with one of the LPS-binding phages. The basal growth rates of the double-phage resistant bacteria, *r_a+b_*, for simultaneous cocktail treatment LUZ19 and PY02, and *r_a+c_*, for cocktail treatment LUZ19 and E215, are calibrated using data set D5 by fitting it with the logistic differential equation (1.6). The fitted curves for these two scenarios are found in Figure 11 (A, B).

**Figure 11:**
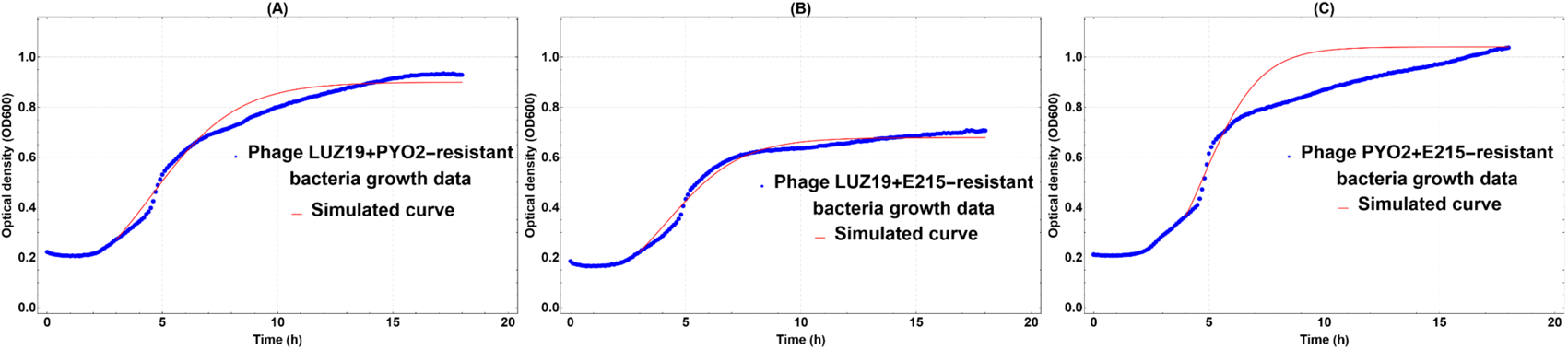
Growth (blue) of P. aeruginosa PA01 resistant to (A) LUZ19+PYO2, (B) LUZ19+E215, or (C) PYO2+E215 over 28 h without the addition of phage. Logistic model fittings (red) to experiment data.

Model (6) is utilized in our simulations when the LPS-binding phages, PYO2 and E215, are applied. Mutational analyses determined that phage resistance to either PYO2 and E215 is conveyed via a mutation at the O-antigen locus of LPS, thereby making the two phages cross-resistant. Therefore, double phage bacterial mutants that emerge from cocktail treatment of PYO2 and E215 harbor a mutation in the *wzy* gene only. Hence, it is reasonable to assume that the basal growth rates of double resistant bacteria that emerge from simultaneous cocktail treatment PYO2 and E215 treatments, *r_b_*_+_*_c_*, and sequential treatments with PYO2 and E215 (*r_b_*_→_*_c_*, *r_c_*_→_*_b_*), as listed in Table 2, are equal to the growth rate of the phage PY02-resistant bacteria, *r_b_*. Indeed, this assumption is corroborated by examining the growth curves between phage PYO2-resistant bacteria (Figure 6B) and cocktail PYO2 and E215-resistant bacteria (Figure 11C). We note that Figure 11C was produced by fitting the logistic differential equation (1.6) to data set D5 (Figure 11C).

By the same reasoning, the mutation rate, *a*, in the two-phage with collateral resistance model (6), is equal to *a_l_*, the mutation rate for LPS-binding phages (See Table 1). The mutation rate *a*_0_*_R_12__*: = *a_lp_* is calibrated using data set D4 and the growth rates described above. The values for all the new parameters associated with models (5) and (6) are listed in Table 2.

As shown in Figure 10, our calibrated two-phage models successfully incorporate parameters from the single phage model to accurately predict the change of the total bacterial population density over time. Both the *in vitro* and *in silico* data found that cocktail treatment LUZ19 and PYO2 is more effective than cocktail LUZ19 and E215 in controlling the total bacterial population density (Figure 10A-B). On the other hand, we observe that cocktail treatment PYO2 and E215 is as effective as single PYO2 treatment (Figure 5B versus Figure 10C). This is consistent with our model for two-phage treatment with collateral resistance, where treatment of two phages is near-synonymous to treatment by a single phage. Thus, the exponential growth of the cocktail PYO2 and E215 resistant bacteria happens much earlier than cocktail LUZ19 and PYO2 or LUZ19 and E215 resistant bacteria because only one SNP is required to gain resistance to both phages.

### 3.7 Fitness cost

We determine the fitness cost of *P. aeruginosa* PAO1 acquiring mutations that allow for phage resistance. Figure 12 illustrates the relative growth rates of *P. aeruginosa* PAO1 with different phage resistance profiles compared to ancestral bacteria. As observed, the single mutants only have a slight decrease in their growth rates relative to the ancestral bacteria. In contrast, the double-mutant strains from cocktail treatment LUZ19 and PYO2 and cocktail treatment LUZ19 and E215 bear significantly higher fitness costs than the single mutants. Among the single mutant strains, the phage strain LUZ19-resistant bacteria has the lowest relative growth rate: 0.88. Whereas phage strain PYO2-resistant bacteria and phage strain E125-resistant bacteria have relatively higher growth rates: 0.91 and 0.92 respectively. Because modifications to conserved components (e.g. LPS) require a fitness trade-off (47), it follows that the same mutation in the *wzy* gene conveys similar reduced growth rates for both the PYO2 and E215 resistant strains. Note that we define “double-mutant strain” as a bacterial strain with mutations in both the pili and LPS loci. The double-mutant strain in the cocktail treatment LUZ19 and PYO2 has a slightly higher fitness cost than the double-mutant strain in the cocktail treatment LUZ19 and E215.

**Figure 12:**
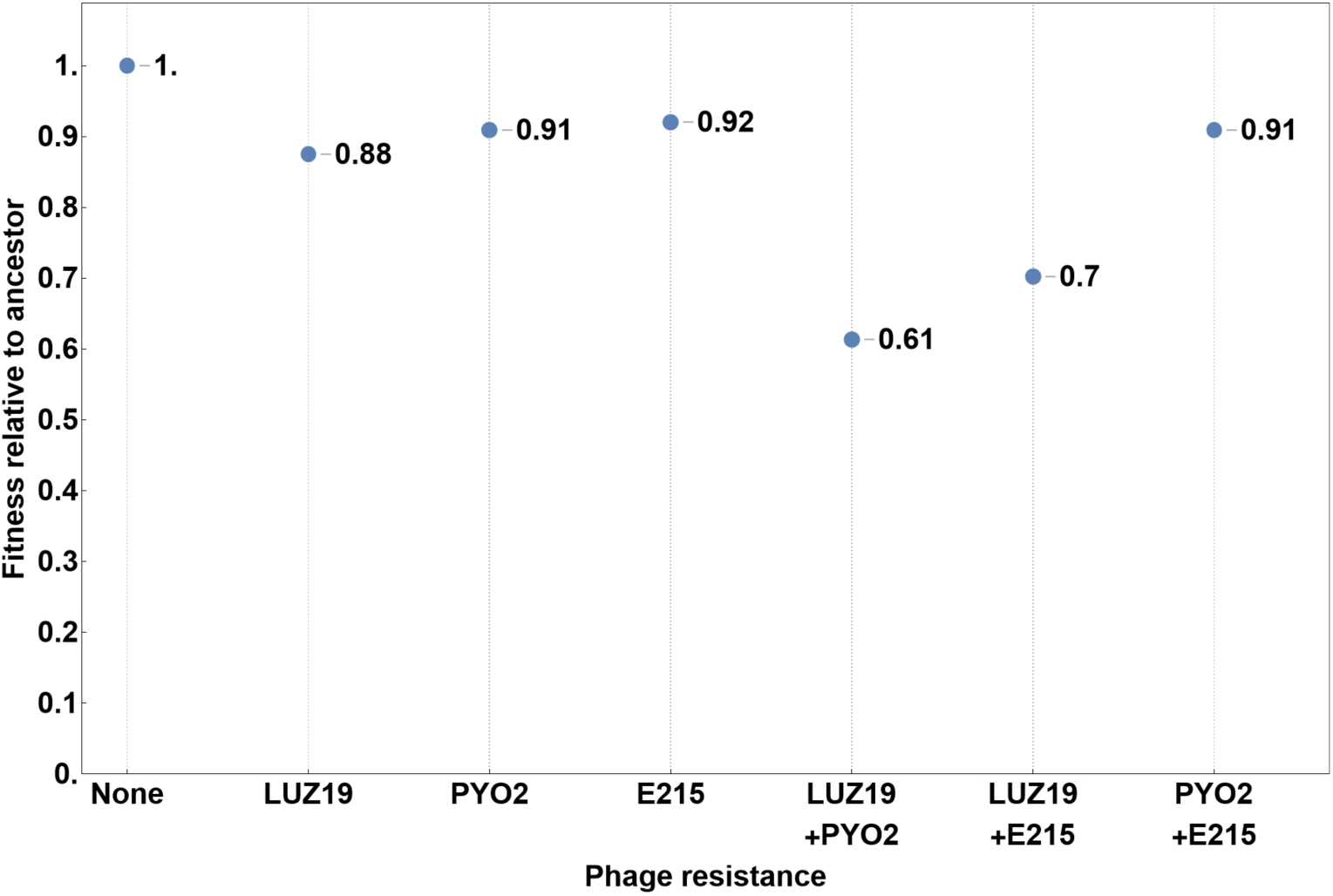
Relative growth rates of different variants of resistant bacteria with respect to the ancestral bacteria.

### 3.8 Sequential phage treatments

Next, we conducted *in silico* experiments of sequential treatment of two phage strains. There are six possible combinations with PYO2, E215, and LUZ19. In simultaneous treatment experiments, a two-phage cocktail (MOI 0.5 per phage, total MOI 1) was added at 0h. To capture the effect of total phage versus total bacteria at 0h and highlight the true effect of sequential treatment, we simulate that the first phage strain is added at MOI 1 at 0h. The second dose is added at 8h at MOI 1. Wright et al. previously suggested that sequential treatments in which the first added phage is pili-binding and the second added phage is LPS-binding can cause significantly lower fitnesses in the double-mutant strains compared to the cocktails with the same phages and the sequential treatments with the same phages applied in a reversed order (11). Therefore, when simulating sequential treatment, we reduced the basal growth rates of the double-mutant strains that emerge in the treatment of LUZ19→PYO2 and LUZ19 → E215, denoted as *r_a_*_→_*_b_* and *r_a_*_→*c*_ respectively, by 10% relative to *r_a_*_)_*_b_* and *r_a_*_)*c*_. Whereas the double-mutant strains that emerge in the treatment of phage PYO2→LUZ19 and E215→LUZ19 are assumed to have the same growth rate as the double-mutant strains in the corresponding simultaneous treatments. That is, *r_b_*_→_*_a_* = *r_a+b_* and *r_c_*_→*a*_ = *r_a+c_*. However, we note that the calibration and assumptions really depend on the strain of phages being applied and hence, the model being used. For this, we remind the reader that phage LUZ19 is a pili-binding phage and phages PYO2 and E215 are LPS-binding. The parameters are listed in Table 2.

To simulate sequential treatments with both pili and LPS binding phages (Figure 13A-D), we utilize the two-phage mathematical model without collateral resistance (Equation (1.4)). As shown in Figure 13A and Figure 13B, both sequential treatment LUZ19→PYO2 and sequential treatment PYO2→LUZ19 cause significant reduction in bacterial density. The sequential treatment LUZ19→PYO2 is more effective than the sequential treatment PYO2→LUZ19 in delaying the exponential growth of the double mutants, indicating the importance of the order of phage addition in sequential treatment. These trends also occur between sequential treatment LUZ19→E215 and sequential treatment E215→LUZ19, as shown in Figure 13C and Figure 13D.

**Figure 13:**
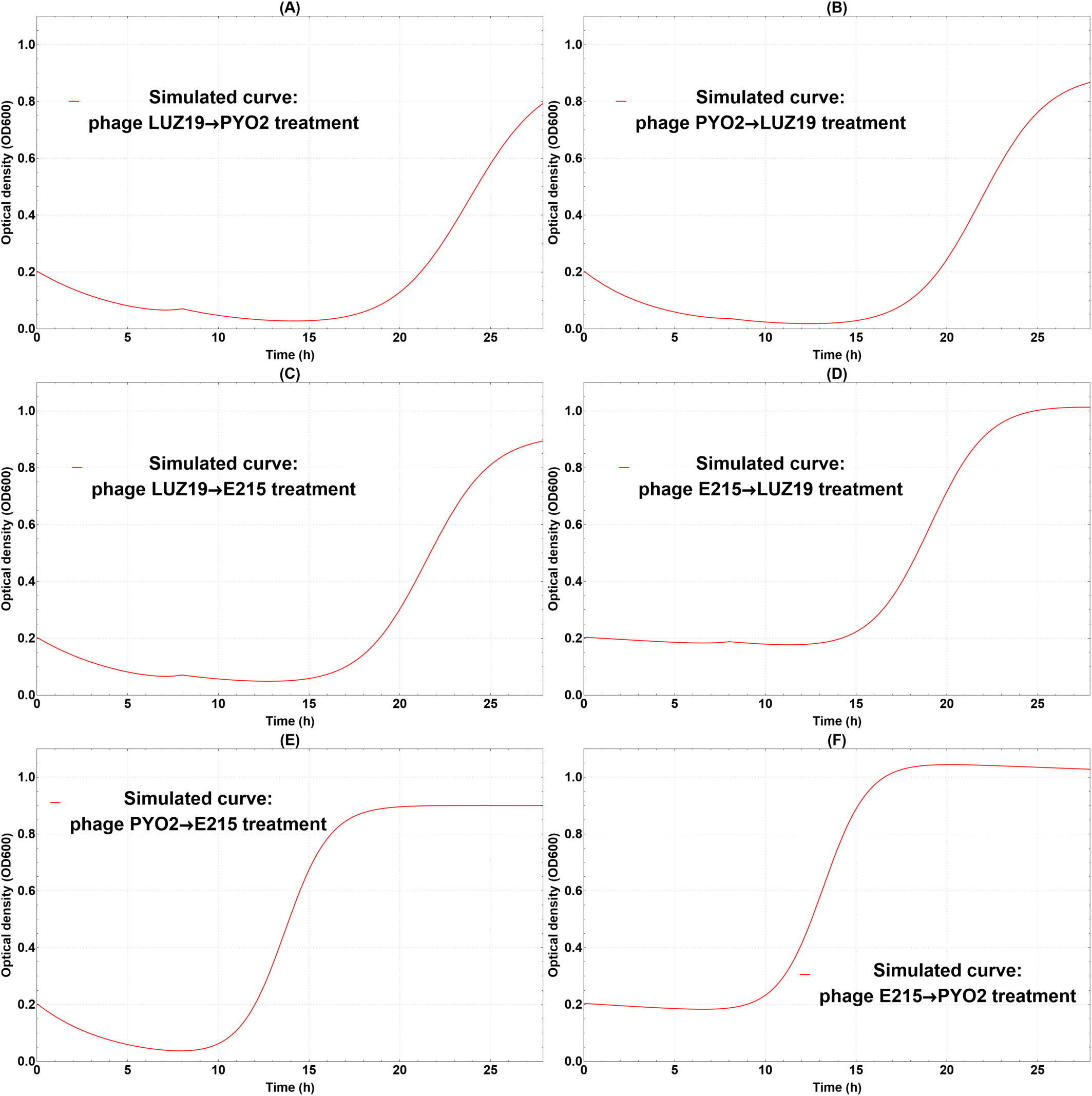
*In silico* experiments for two-phage sequential treatments. Each curve simulates the total bacteria density after adding one phage strain at MOI 1 at 0h, and a different phage strain at MOI 1 at 8h. (A)LUZ19→PYO2; (B) PYO2→LUZ19; (C) LUZ19→E215; (D) E21 →LUZ19; (E) PYO2→E215; (F) E215→PYO2

When simulating sequential treatment with two LPS binding phages, we refer to the two-phage mathematical model with collateral resistance (Equation (1.5)). Consistent with trends observed *in vitro*, sequential treatment PYO2→E215 and sequential treatment E215→PYO2 perform similar to the two-phage cocktail treatment PYO2 and E215 in delaying the exponential growth of LPS-resistant mutants, as shown in Figure 13E and Figure 13F.

Our simulations suggest that all six sequential treatments are less effective at controlling the total bacteria population density than their two-phage simultaneous treatment counterparts (Figure 13 vs Figure 10).

### 3.9 Model solution: Two-phage models

For the double phage model with collateral resistance (Figure 14C, F, and I), the model starts with ancestral bacteria and two phage strains that both target the same receptor on bacteria. The resulting bacterial growth transients in this model are similar to those from the single phage model (Figure 7) of PYO2 or E215, except that both of the LPS-binding phages are present in the system.

**Figure 14:**
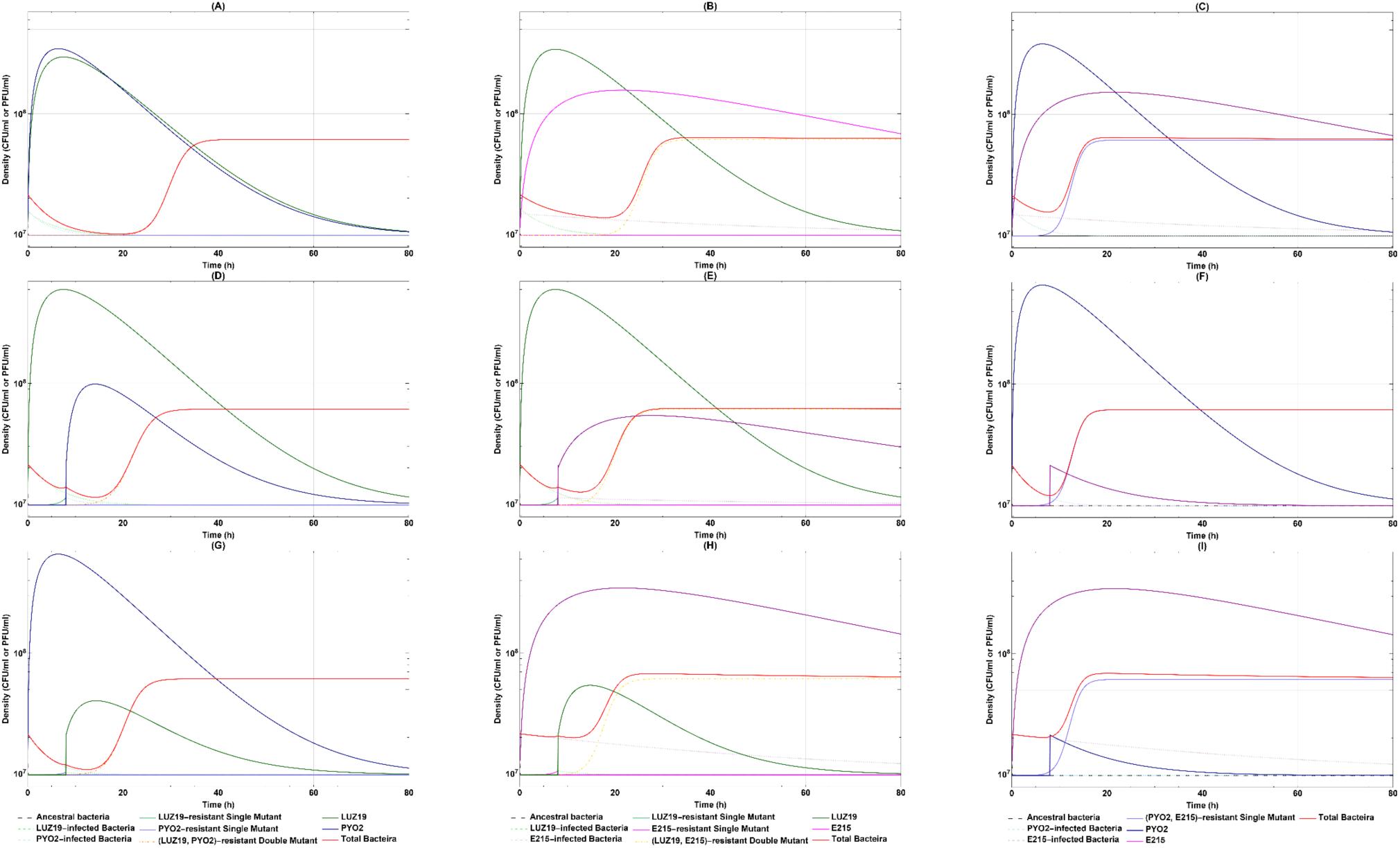
Double phage model numerical simulations. For all the simulations, the black dashed line denotes the ancestral bacteria, and the red solid line denotes all the bacteria. The infected bacterial strains are denoted by dashed lines with different colors. The single mutant strains and the phage strains are denoted by solid lines with different colors. The double mutant strains are denoted by dot dashed lines with different colors. (A-C) Double phage simultaneous treatments with both phage strains are added at time 0 h: the MOI of each phage strain is 0.5. (D-I) Double phage sequential treatments with first dose at time 0h and second dose at time 8h: the MOI of each phage strain is 1. (Column 1) LUZ19 and PYO2: no collateral resistance. (Column 2) LUZ19 and E215: no collateral resistance. (Column 3) PYO2 and E215: collateral resistance involved. All phage specific and treatment specific parameters are listed in Table 1 and 2. For all the simulations, *B*(*0*) = *0*.*2*, and *K* = *0.*9. Note that the OD_600_ is converted to CFU/mL by using y = 57115528.37x + 9785470.84, where x is the OD_600_ value.

In the absence of collateral resistance, we have two scenarios that better delay resistance: phages added simultaneously and sequentially. For the simultaneous treatment (Figure 14 A and B), we start with ancestral bacteria and two different phage strains. Similar to the predicted single phage model, the ancestral bacteria population is wiped out quickly, and the populations of both phage strains increase significantly. Eventually, the double mutant becomes dominant. As with the sequential phage treatment (Figure 14 D, E, G, and H), the outcome of the sequential phage treatment (pili → LPS phage order) is dominated by the double resistant mutant. However, the double mutant arises much earlier for the simulated sequential phage treatment than the simultaneous phage treatment.

### 3.10 Exploring second dose timing

As previously mentioned, a limitation to performing *in vitro* characterization of different phage-bacteria combinations is the variety of ways in which the system can be manipulated. Phage order, dose, and timing are all variables that impact the timing and onset of phage resistance by bacteria. In order to explore how second dose timing will affect the double phage sequential treatments in terms of their ability to control total bacterial population density and minimize the resistance, we extended the previous sequential model to include more possible second dose timings and quantified the abilities to control total bacterial population density and minimize the resistance by measuring the total bacteria density (Figure 15A) and the proportion of the bacteria that have complete resistance to that treatment (Figure 15B), at 15h. Similar to other antimicrobial agents, the success of phage therapy depends on both its ability to suppress the bacterial populations and limit the emergence of phage resistance. Certain phages may be effective at clearing a sensitive bacterial population, but the remaining bacterial population contains a high proportion of bacteria that are resistant to all phages used in those treatments, which may potentially lead to extensive bacterial population regrowth. In Figure 15A, we compared different double phage sequential treatments based on their ability to control the total bacterial population density. In particular, we used the total bacterial population density after 15h as an indicator value, as shown by the y-axis in Figure 15. We used the same carrying capacity *K* = *0.*9 while simulating the model for every treatment such that at 15h, the total bacterial population density will not reach the carrying capacity. The initial bacterial density for all the simulations in this section is 0.2, and the initial phage density for all the simulations is the same as the initial bacterial density (MOI 1).

**Figure 15:**
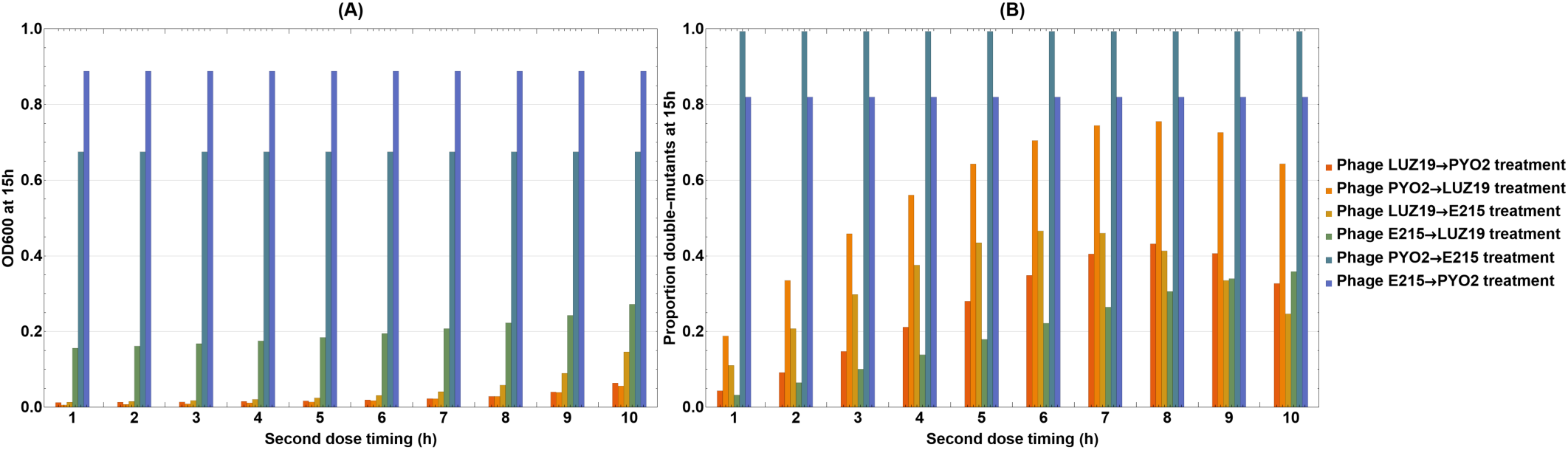
Comparison of double phage sequential treatments with different phage combinations, phage orders, and second dose timings in terms of their ability to control total bacterial population density and minimize the resistance: (A) The total bacterial density at 15h (B) The proportion of the bacteria that are resistant to all phages used in that treatment at 15h. Note that different colors indicate different phage combinations and orders.

In general, as second dose timing increases, the treatments become less effective in controlling the total bacterial population density. Treatments with collateral resistance are insensitive to a change in second dose timings. Moreover, sequential LUZ19 → PYO2 and PYO2 → LUZ19 are the two most effective phage sequences in suppressing bacterial growth. LUZ19 → E215 sequential treatment is slightly less effective than LUZ19 → PYO2, but LUZ19 → E215 sequential treatment’s efficacy of controlling the total bacterial population density is more sensitive to the change in its second dose timing than the phage (LUZ19 → PYO2) sequential treatment.

In Figure 15B we compared the same set of sequential treatments based on their ability to minimize the proportion of phage resistant bacteria. For all the sequential treatments without collateral resistance, as second dose timing increases, the ability of minimizing resistance decreases then increases.

Increasing second dose timing elongates growth period for the single mutants, and increase the mutation supply of the double mutants from single mutants through one-step evolution. Thus, an initial decrease in the ability of minimizing resistance is observed. However, as the second dose timing becomes large enough, the growth of the single mutants is going to suppress the growth of the double mutants, and this will reduce the proportion of the double mutants. Among all the sequential treatments without collateral resistance, the E215→LUZ19 sequential treatment is the most effective one in minimizing the resistance, though it is not the best one in controlling the total bacterial density. Phage LUZ19 → PYO2 and LUZ19 → E215 are the next two most effective sequential treatments in minimizing the resistance, and they are also efficient in controlling the total bacterial density. Thus, we conclude that LUZ19 → PYO2 and LUZ19→E215 sequential treatments are the two most successful treatments. PYO2→E215 and E215→PYO2 sequential treatments’ abilities of minimizing resistance are not sensitive to the second dose timing because the bacteria that are resistant to all phage strains used in these two treatments are actually the single mutants that exhibit collateral resistance to both phage PYO2 and E215.

### 3.11 Decreased binding rate and different phage combinations

Next, we determined the effects of decreasing the phages adsorption rate to simulate greater competition between different sensitive host strains. Furthermore, we used different phage combinations in these two scenarios in order to study the competition between two phages on one host bacteria. As shown in Figure 16A, we have a phage with low potency (low lysis rate) and high burst size and a phage with high potency and low burst size. Initially, the phage with higher potency proliferates faster than the other phage, but its population shrinks quickly after 23h when there is no longer enough susceptible bacteria and phage reservoir (infected bacteria). In contrast, the phage with low potency and high burst size can reach a much higher maximum population density due to its large burst size. In addition, due to low binding rate, ancestral bacteria can exist for a much longer period of time, resulting in the delayed exponential growth of the single mutants due to the competition between single mutants and ancestral bacteria. As ancestral bacteria die out, the single mutants become dominant, which delays the exponential growth of the double mutants due to competition. Thus, although decreased binding rate reduces the ability of phage to proliferate and kill susceptible bacteria, it promotes competition between different bacterial strains and thereby delays the occurrence of multi-phage resistant bacteria. In Figure 16B, we model a phage with high potency and high burst size and a phage with low potency and low burst size. We found that the phage with high potency and high burst size replicates faster and its population dominates over the other phage. Subsequently, single mutants that are resistant to the phage with high potency and high burst size dominate over the other bacterial strains for a long period of time, and the growth of the double mutants is greatly obstructed by these single mutants.

**Figure 16:**
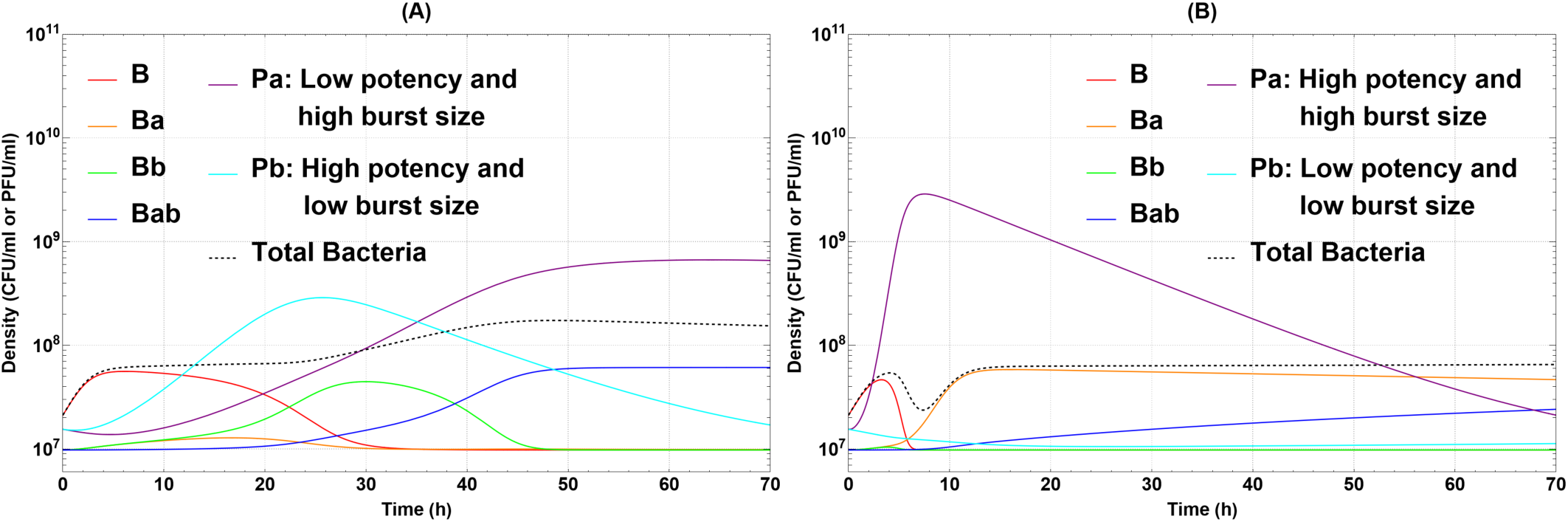
Comparison of two different double phage simultaneous treatments under decreased phage binding rate b: (A) Phage strain with low lysis rate s and high burst size h coupled with the other strain with high potency and low burst size; (B) Phage strain with high potency and high burst size coupled with the other strain with low potency and low burst size. The OD_600_ is converted to CFU/mL by using y = 57115528.37x + 9785470.84, where x is the OD_600_ value.

## 4 Discussion

In this study, we demonstrated experimentally and mathematically that the simultaneous administration of select phage strains as a cocktail was more effective at controlling *P. aeruginosa* density than the phages applied sequentially or individually. By combining *in vitro* and *in silico* models of time-kill kinetics, we show 68 permutations of single, double simultaneous, or double sequential phage treatments. Of these, we found that double simultaneous treatment with LUZ19 and PYO2 was the most potent at reducing bacterial density. The increased potency also promoted a more robust phage growth of both LUZ19 and PYO2 over a longer period of time. In contrast, double simultaneous treatment with different cocktails of LUZ19 and E215 or PYO2 and E215 exhibited weaker potency. In addition, double sequential treatments that initially lack compounding phage infectivities were also lower in potency. Sequential treatment potency could be increased comparable to double simultaneous LUZ19 and PYO2 treatment by reducing the interval between administrations to less than 2h. As to be expected, we show that combining phages that bind to different cell surface receptors suppressed bacterial numbers far longer than combining phages that bind to the same receptor. Nevertheless, evolution of resistance could not be prevented with a cocktail composed of phages that bind to different cell surface receptors. Resistance did impose fitness costs on emerged mutants by hampering their growth by as much as 40%. Together, a formulation with two highly potent phage strains and those that bind asymmetrically were determined to be essential cofactors for superior treatment efficacy of a phage cocktail.

Lytic activity is a critical quality attribute to phage therapy. Of the three phages, PYO2 was the most potent at reducing bacterial numbers. Whereas E215 was unable to reduce bacteria, rather it could only restrain bacterial growth when administered at the same MOI. Stronger potency was achieved when PYO2 and LUZ19 were combined, leading to further reduction in bacterial density by up to 15%. On the other hand, combining two phages that exhibit a high potency and a low potency provided no greater bacterial suppression than the most potent phage strain in that cocktail. Phage replication is divided into the phases of virion attachment, DNA entry, replication, virion assembly and finally, egress via cell lysis. It is not clear how PYO2 was the most potent at reducing *P. aeruginosa*, nor why combining multiple phages lead to a higher potency of a cocktail. These findings suggest that certain phages have a genetic constitution better suited for exploiting host cell surface receptors, intracellular resources, DNA-synthesis and protein-synthesis systems, and/or degrading structural components (48, 49). For other phage strains, the opposite may be true. Therefore, treatment potency is an indication of not only the quantity required to produce a lytic effect, but also the lysis efficiency of the phage agents. The complexity of each phage strain implies that determining individual phage potency is required for adequate cocktail development.

Moreover, treatments employing two potent phage strains that have asymmetrical cell surface receptors was the strongest predictor of therapy ‘longevity’. Double simultaneous LUZ19 and PYO2 treatment provided the longest suppression of the *P. aeruginosa* population as compared to other treatment scenarios. Cocktail treatment eventually failed due to resistance development to both phage strains. By combining phages with asymmetrical receptors, the joint probability of two or more resistance mutations occurring in a single bacterial cell would be governed by the multiplication rule of probabilities (50). Although two nonsynonymous mutations within a single cell occurred during cocktail treatment, the emergence of a double mutant was significantly delayed due to a lower probability of occurrence of both mutations. That is, the probability of occurrence of both receptor *pil* and *wzy* gene mutations is equal to the product of the probability of *wzy* gene mutation occurring and the conditional probability that event *pil* mutation occurring given that *wzy* gene mutation occurs. This implies that phages in a cocktail can ‘synergize’ to extend treatment longevity by multiplying their resistance probabilities. Next, we investigated whether administering phages sequentially was viable as an alternative strategy to suppress the evolution of phage resistance rather than only delaying resistance (13). Our mathematical model predicts that double sequential phage therapy would not be as effective as double simultaneous therapy at preventing resistance (Figure 15). We theorize that sequential treatment does not fully exploit the multiplication rule of probabilities across the bacterial population. Sequential treatment allows a host cell to rapidly acquire a mutation to escape the first phage and these resistant mutants would grow unabated until the addition of the second phage strain. The secondary phage would then encounter two bacterial subpopulations, one sensitive to both phage strains and the other sensitive to only the second phage. The latter subpopulation would not experience a multiplication rule of probabilities and thus the probability of gaining a resistance mutation would be similar to administering a single phage alone. Therefore, we predict that the appearance of mutants resistant to both phage strains would occur sooner than if the phages were treated simultaneously.

Phage resistance remains a major challenge for phage therapy (5, 51). The fate of evolved resistance mutants however is determined in part by their fitness to compete with non-mutant counterparts and maintain its virulence (52). We found *in vitro* and *in silico* that single receptor gene mutations did not impose fitness trade-offs in growth rate under prevailing conditions in the absence of phage. However, evolved mutants with two receptor gene mutations (e.g., *pil* and *wzy*) experienced a cost to resistance, seen by a decline in growth rate of up to 40% in the absence of phages compared to ancestral cells. These findings agreed with previous reports that single-receptor mutations may be of little consequence to the bacterial cell, whereas multiple mutations elicit an unavoidable cost. Wright et al. similarly showed that the growth rate of evolved independent resistance to two phage strains was significantly decreased (11). Markwitz et al. further identified other fitness costs to evolved resistance to multiple *P. aeruginosa* phage strains including motility, virulence, and sensitivity to human serum (53). This implies that evolved multi-phage resistant cells can outcompete non-mutant sensitive bacteria in the presence of phages, but resistant variants would be at a growth disadvantage in the absence of phages. In the body, this may occur between phage cocktail administrations due to the rapid decay of phages (10, 54).

Phages are unlike other antimicrobials in that they self-dose (i.e. replicate) during treatment. When PYO2 and LUZ19 are administered together, our mathematical model predicts that both phages would grow evenly to a similar maximum density. In contrast, administering either PYO2 and E215 or LUZ19 and E215 together, the phages grow unevenly. E215 is predicted to replicate not as well nor reach as high concentration when paired with either PYO2 or LUZ19. Since a bacterial cell can only be infected by a single phage, increasing phage strain diversity also introduces the possibility for phage competition (20). As mentioned, host competitive ability varied among the three phages tested. The reduction in productivity that E215 may experience in the presence of competitors provides evidence that multiple infections can have a severe impact on self-dosing. Because host competitive ability was independent of pairing phages that use the same cell surface receptor (PYO2 and E215), phage compete in replication efficiency. E215 replication cycle takes longer than the other phages and thus, there is a lag of new progeny to infect susceptible cells. Susceptible cells are monopolized by the more efficiently replicating phages, further perpetuating E215 growth demise as its progeny have fewer new cells to infect. This competitive exclusion suggests two phages cannot coexist when infecting the same host bacterium. Betts et al. similarly found that pairwise phages in double simultaneous treatments experienced less viral growth in the absence of phage competition (20). We found further evidence of this when simulating sequential treatments. Our mathematical model predicts that higher total phage growth occurred during double simultaneous treatment compared to double sequential treatment. This is because all susceptible cells initially supported single phage replication instead of supporting replication of two phages. By the time the second phage strain is administered, the susceptible population numbers are low and unable to support robust amplification of the second phage. In addition, both phages are now competing for the small population of susceptible cells. This would suggest that simultaneous treatment provides greater productivity of each phage component to maximize self-dosing. The genetic and physiological factors that determine the outcome of host competition appear to be of critical importance to cocktail design, yet they remain poorly understood.

Mathematical tools are instrumental to understanding complex biological systems. Models help us parameterize, control, and optimize our predictions while allowing us to study the effects of changing each component and/or treatment condition in the system. In this study the use of our mathematical models allowed us to predict the growth of the phage population which was undetectable using spectrophotometry. Furthermore, we parameterized bacterial growth characteristics to predict bacterial density under the experimental limit of detection. Visualizing the change in individual phage strain and bacterial abundance through model simulations helped us understand the previously discussed bacteria-phage kinetics. Our mathematical models incorporated both the phage-phage synergy and the stepwise evolution of the bacteria to provide a mechanistic description of the kinetics between the bacteria and two phage strains, extending the model work of Payne & Jansen and Cairns (27, 28). In addition, we have introduced two versions of the double phage model, with and without collateral resistance. Thus, when novel phages with known receptors are introduced our models can use individual phage killing and evolutionary parameters to infer the treatment’s efficacy. We calibrated our models and used them as plug-and-play frameworks to explore more treatment regimens and scenarios, such as the sequential treatments with different dosing intervals and solutions with decreased phage binding rate and hypothetical phage choices. Unlike previous studies that mainly focused on the phage killing aspect of phage therapy, our double phage models incorporate both phage killing and the evolution of bacterial phage resistance (26, 55, 56). Understanding both is important for the development of combination phage therapy in the future.

In conclusion, we found that the optimal pairwise phage treatment strategy was the double simultaneous administration of two highly potent and asymmetrically binding phage strains. This treatment regimen had the greatest lysis efficiency, reduced bacterial density the most, and suppressed the evolution of resistance for the longest duration. However, this cocktail treatment strategy could not prevent phage resistance *in vitro*. Further study of three phage cocktails may limit resistance since the multiplication of two mutation probabilities only led to delayed resistance. We found that double sequential phage treatment had several drawbacks compared to double simultaneous administration. However, a sequential regimen may have other benefits not explored in this study, such as limiting the emergence of neutralizing anti-phage immune responses (57). Together, these results highlight that the pharmacokinetics and pharmacodynamics of each phage are important in designing a therapeutic cocktail. Future work to map which phage properties enhance potency is necessary to optimize multi-phage cocktail treatment regimens.

## Acknowledgments

The authors thank the American Institute of Mathematics for providing support through its Structured Quartet Research Ensembles program. This research group was initially formed during the Collaborative Workshop for Women in Mathematical Biology hosted by the Institute for Pure and Applied Mathematics at the University of California, Los Angeles in June 2019. The authors would like to thank Dr. Aadrita Nandi for her discussion.

## Appendix A: Local sensitivity analysis

The model needs to start at an admissible point in parameter space so we performed modeling fitting first. Then we use the model transients to generate the discretized sensitivity matrix S. We then used S to rank parameters by sensitivity and set a threshold such that parameters with sensitivity below the threshold (insensitive) are fixed and parameters with sensitivity above the threshold (sensitive) are explored. All four observable model outputs (*B*, *B_I_*, *B_R_* and *P*) were sampled at 25 time points (each hour for 25 hours). Given that there are 9 model parameters explored, a 100 × 9 discretized sensitivity matrix S is produced.

Next, we ranked the impact of each parameter on all four observable model outputs (*B*, *B_I_*, *B_R_* and *P*) by calculating a root mean square sensitivity measure, as defined in Brun et al. (58). For each column j of the normalized sensitivity matrix, we get

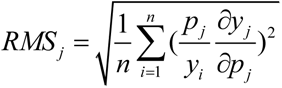

Parameter j is deemed insensitive if RMS_j_ is less than 5% of the value of the maximum RMS value calculated over all parameters. By this measure, the only parameter deemed sensitive for all three phage strains is *s*, as shown in Figure 8. It is interesting to note that phage PYO2 and phage E215 also showed high sensitivity to *h*, *r_r_*, and *b*. Phage A showed sensitivity to *r_n_*. Again, the mechanism of infection for phage LUZ19 differs from that of phage PYO2 and E215 and that difference is evident in the system activity.

## Supplementary Materials

**Supplementary Table 1:** Genotypic and phenotypic characteristics of phage strains.

| Phage name | Genome (bp) | # ORFs | Accession # | K<sub>adsorption</sub> (ml/min) | Eclipse period (min) | Latent period (min) | Burst size (PFU/cell) |
| --- | --- | --- | --- | --- | --- | --- | --- |
| LUZ19<sup>a</sup> | 43,548 | 54 | NC_010326.1 | $1.2 \times 10^{-7}$ | ND | 22 | 122 |
| PYO2<sup>b</sup> | 72,697 | 91 | MF490236.1 | $1.8 \times 10^{-10}$ | 24 | 26 | 183 |
| E215<sup>b</sup> | 66,789 | 95 | MF490241.1 | $1.4 \times 10^{-9}$ | 26 | 30 | 195 |

<sup>a</sup> Growth characteristics were determined by Ceysens, PJ. et al. 2011.
<sup>b</sup> Growth characteristics were determined in this study on *P. aeruginosa* PAO1.

**Supplementary Figure 1:**
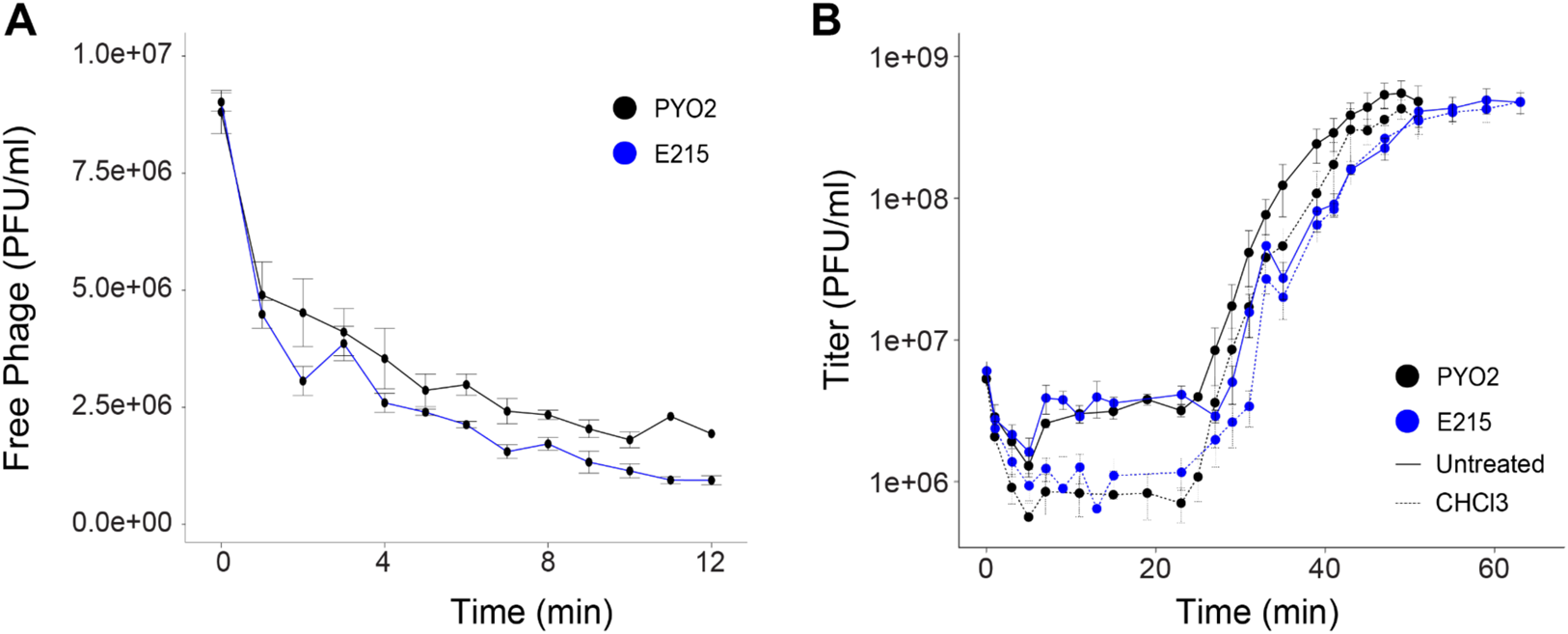
PYO2 (black) and E215 (blue) growth characteristics during incubation at 37°C with shaking at 120rpm. (A) Adsorption rates (K, phage^−1^ cell^−1^ mL^−1^ min^−1^) were determined by measuring the decline of free-phage in a bacteria-phage solution at a MOI of 0.02. Measurements were taken every minute for 12 min. Adsorption rates (K_adsorption_) were calculated when 50% of free-phage remained in solution. **(B)** One-step growth curves were performed to determine the eclipse, latent period, and burst size per cell of phage infection. Phages were mixed with bacteria at an MOI of 0.1 and samples were taken every 2 minutes until the plateau of free-phage. Samples were divided to calculate the eclipse period (derived from CHCl3 treatment), latent period (derived from untreated samples), and burst size (calculated from the average plateau point between untreated and CHCl3 treatment divided by the difference in titer at t=7 of untreated and treated samples).

**Supplementary Table 2:** Identity, function, position, and types of mutations identified *in vitro*.

| Phage treatment | # CFUs | Gene name | Gene function | Gene description | Chromosome position | BRESEQ annotation |
| --- | --- | --- | --- | --- | --- | --- |
| PYO2 | 3 | wzy | LPS biosynthesis | Oligosaccharide repeat unit polymerase | 2694486 | insertion |
| E215 | 1 | wzy |  | Oligosaccharide repeat unit polymerase | 2694486 | insertion |
| E215 | 1 | wzy |  | Oligosaccharide repeat unit polymerase | 2694975 | insertion |
| PYO2 + E215 | 2 | wzy |  | Oligosaccharide repeat unit polymerase | 2694486 | insertion |
| PYO2 + E215 | 1 | wzy |  | Oligosaccharide repeat unit polymerase | 2695229 | deletion |
| LUZ19 + PYO2 | 1 | wzy |  | Oligosaccharide repeat unit polymerase | 2694486 | insertion |
| LUZ19 + E215 | 1 | wzy |  | Oligosaccharide repeat unit polymerase | 2695221 | missense |
| LUZ19 + E215 | 1 | wzy |  | Oligosaccharide repeat unit polymerase | 2694486 | insertion |
| LUZ19 + E215 | 1 | wzy |  | Oligosaccharide repeat unit polymerase | 2694975 | insertion |
| LUZ19 | 1 | pilF | Type IVa pilus system | Type IV pilus biogenesis protein | 1970808 | nonsense |
| LUZ19 | 1 | pilR |  | Type IV fimbriae expression regulatory protein | 1203506 | insertion |
| LUZ19 | 1 | pilB |  | Type IV fimbrial assembly, ATPase | 1228607 | missense |
| LUZ19 + PYO2 | 1 | pilM |  | Type IV pilus biogenesis protein | 619280 | nonsense |
| LUZ19 + E215 | 1 | pilQ |  | Type IV pilus biogenesis protein | 623834 | deletion |
| LUZ19 + E215 | 1 | pilN |  | Type IV pilus biogenesis protein | 620560 | nonstop |
| LUZ19 + E215 | 1 | pilO |  | Type IV pilus biogenesis protein | 620560 | missense |
| LUZ19 + E215 | 1 | pilR |  | Type IV pilus biogenesis protein | 1203161 | missense |

## References

1. Sader HS, Castanheira M, Duncan LR, Flamm RK. Antimicrobial susceptibility of Enterobacteriaceae and Pseudomonas aeruginosa isolates from United States medical centers stratified by infection type: results from the International Network for Optimal Resistance Monitoring (INFORM) surveillance program, 2015-2016. Diagnostic Microbiology and Infectious Disease. 2018;92(1):69–74.

2. Tabak YP, Merchant S, Ye G, Vankeepuram L, Gupta V, Kurtz SG, et al. Incremental clinical and economic burden of suspected respiratory infections due to multi-drug-resistant Pseudomonas aeruginosa in the United States. Journal of Hospital Infection. 2019;103(2):134–41.

3. Simpkin VL, Renwick MJ, Kelly R, Mossialos E. Incentivising innovation in antibiotic drug discovery and development: progress, challenges and next steps. Journal of Antibiotics (Tokyo). 2017;70(12):1087–96.

4. Doster E, Rovira P, Noyes NR, Burgess BA, Yang X, Weinroth MD, et al. Investigating effects of tulathromycin metaphylaxis on the fecal resistome and microbiome of commercial feedlot cattle early in the feeding period. Frontiers in Microbiology. 2018;9:1715.

5. Luong T, Salabarria AC, Roach DR. Phage therapy in the resistance era: where do we stand and where are we going? Clinical Therapeutics. 2020;42(9):1659–80.

6. Hatfull GF, Dedrick RM, Schooley RT. Phage therapy for antibiotic-resistant bacterial infections. Annual Review of Medicine. 2022;73:197–211.

7. Chan BK, Stanley G, Modak M, Koff JL, Turner PE. Bacteriophage therapy for infections in CF. Pediatric Pulmonology. 2021;56 Suppl 1:S4–S9.

8. Chan BK, Sistrom M, Wertz JE, Kortright KE, Narayan D, Turner PE. Phage selection restores antibiotic sensitivity in MDR Pseudomonas aeruginosa. Scientific Reports. 2016;6:26717.

9. Forti F, Roach DR, Cafora M, Pasini ME, Horner DS, Fiscarelli EV, et al. Design of a broad-range bacteriophage cocktail that reduces *Pseudomonas aeruginosa* biofilms and treats acute infections in two animal models. Antimicrobial Agents and Chemotherapy. 2018;62(6).

10. Roach DR, Leung CY, Henry M, Morello E, Singh D, Di Santo JP, et al. Synergy between the host immune system and bacteriophage is essential for successful phage therapy against an acute respiratory pathogen. Cell Host Microbe. 2017;22(1):38–47 e4.

11. Wright RCT, Friman VP, Smith MCM, Brockhurst MA. Resistance evolution against phage combinations depends on the timing and order of exposure. mBio. 2019;10(5).

12. Yang Y, Shen W, Zhong Q, Chen Q, He X, Baker JL, et al. Development of a bacteriophage cocktail to constrain the emergence of phage-resistant *Pseudomonas aeruginosa*. Frontiers in Microbiology. 2020;11:327.

13. Hall AR, De Vos D, Friman VP, Pirnay JP, Buckling A. Effects of sequential and simultaneous applications of bacteriophages on populations of Pseudomonas aeruginosa in vitro and in wax moth larvae. Applied and Environmental Microbiology. 2012;78(16):5646–52.

14. Law N, Logan C, Yung G, Furr CL, Lehman SM, Morales S, et al. Successful adjunctive use of bacteriophage therapy for treatment of multidrug-resistant Pseudomonas aeruginosa infection in a cystic fibrosis patient. Infection. 2019;47(4):665–8.

15. Chan BK, Turner PE, Kim S, Mojibian HR, Elefteriades JA, Narayan D. Phage treatment of an aortic graft infected with Pseudomonas aeruginosa. Evolution, Medicine, and Public Health. 2018;2018(1):60–6.

16. Aslam S, Courtwright AM, Koval C, Lehman SM, Morales S, Furr CL, et al. Early clinical experience of bacteriophage therapy in 3 lung transplant recipients. American Journal of Transplantation. 2019;19(9):2631–9.

17. Oechslin F. Resistance development to bacteriophages occurring during bacteriophage therapy. Viruses. 2018;10(7).

18. Dedrick RM, Smith BE, Garlena RA, Russell DA, Aull HG, Mahalingam V, et al. *Mycobacterium abscessus* strain morphotype determines phage susceptibility, the repertoire of therapeutically useful phages, and phage resistance. mBio. 2021;12(2).

19. Mutalik VK, Adler BA, Rishi HS, Piya D, Zhong C, Koskella B, et al. High-throughput mapping of the phage resistance landscape in E. coli. PLOS Biology. 2020;18(10):e3000877.

20. Betts A, Gifford DR, MacLean RC, King KC. Parasite diversity drives rapid host dynamics and evolution of resistance in a bacteria-phage system. Evolution. 2016;70(5):969–78.

21. O’Flynn G, Ross RP, Fitzgerald GF, Coffey A. Evaluation of a cocktail of three bacteriophages for biocontrol of *Escherichia coli* O157:H7. Applied and Environmental Microbiology. 2004;70(6):3417–24.

22. Summers AO. Generally overlooked fundamentals of bacterial genetics and ecology. Clinical Infectious Diseases. 2002;34 Suppl 3:S85–92.

23. Abedon ST, Thomas-Abedon C. Phage therapy pharmacology. Current Pharmaceutical Biotechnology. 2010;11(1):28–47.

24. Delattre R, Seurat J, Haddad F, Nguyen TT, Gaborieau B, Kane R, et al. Combination of in vivo phage therapy data with in silico model highlights key parameters for pneumonia treatment efficacy. Cell Reports. 2022;39(7):110825.

25. Rodriguez-Gonzalez RA, Leung CY, Chan BK, Turner PE, Weitz JS. Quantitative models of phage-antibiotic combination therapy. mSystems. 2020;5(1).

26. Schmerer M, Molineux IJ, Bull JJ. Synergy as a rationale for phage therapy using phage cocktails. PeerJ. 2014;2:e590.

27. Payne RJ, Jansen VA. Understanding bacteriophage therapy as a density-dependent kinetic process. Journal of Theoretical Biology. 2001;208(1):37–48.

28. Cairns BJ, Timms AR, Jansen VA, Connerton IF, Payne RJ. Quantitative models of in vitro bacteriophage-host dynamics and their application to phage therapy. PLOS Pathogens. 2009;5(1):e1000253.

29. Xie Y, Wahab L, Gill JJ. Development and validation of a microtiter plate-based assay for determination of bacteriophage host range and virulence. Viruses. 2018;10(4).

30. Storms ZJ, Teel MR, Mercurio K, Sauvageau D. The virulence index: a metric for quantitative analysis of phage virulence. Phage (New Rochelle). 2020;1(1):27–36.

31. Blanco-Cabra N, Lopez-Martinez MJ, Arevalo-Jaimes BV, Martin-Gomez MT, Samitier J, Torrents E. A new BiofilmChip device for testing biofilm formation and antibiotic susceptibility. NPJ Biofilms Microbiomes. 2021;7(1):62.

32. Chin WH, Kett C, Cooper O, Museler D, Zhang Y, Bamert RS, et al. Bacteriophages evolve enhanced persistence to a mucosal surface. Proceedings of the National Academy of Sciences of the United States of America. 2022;119(27):e2116197119.

33. Luong T, Salabarria AC, Edwards RA, Roach DR. Standardized bacteriophage purification for personalized phage therapy. Nature Protocols. 2020;15(9):2867–90.

34. Lammens E, Ceyssens PJ, Voet M, Hertveldt K, Lavigne R, Volckaert G. Representational difference analysis (RDA) of bacteriophage genomes. Journal of Microbiological Methods. 2009;77(2):207–13.

35. Ceyssens PJ, Glonti T, Kropinski NM, Lavigne R, Chanishvili N, Kulakov L, et al. Phenotypic and genotypic variations within a single bacteriophage species. Virology Journal. 2011;8:134.

36. Kropinski AM. Measurement of the rate of attachment of bacteriophage to cells. Methods in Molecular Biology. 2009;501:151–5.

37. Chevallereau A, Blasdel BG, De Smet J, Monot M, Zimmermann M, Kogadeeva M, et al. Next-generation “-omics” approaches reveal a massive alteration of host RNA metabolism during bacteriophage infection of *Pseudomonas aeruginosa*. PLOS Genetics. 2016;12(7):e1006134.

38. Chen S, Zhou Y, Chen Y, Gu J. fastp: an ultra-fast all-in-one FASTQ preprocessor. Bioinformatics. 2018;34(17):i884–i90.

39. Bankevich A, Nurk S, Antipov D, Gurevich AA, Dvorkin M, Kulikov AS, et al. SPAdes: a new genome assembly algorithm and its applications to single-cell sequencing. Journal of Computational Biology. 2012;19(5):455–77.

40. Bosi E, Donati B, Galardini M, Brunetti S, Sagot MF, Lio P, et al. MeDuSa: a multi-draft based scaffolder. Bioinformatics. 2015;31(15):2443–51.

41. Aziz RK, Bartels D, Best AA, DeJongh M, Disz T, Edwards RA, et al. The RAST Server: rapid annotations using subsystems technology. BMC Genomics. 2008;9:75.

42. Deatherage DE, Barrick JE. Identification of mutations in laboratory-evolved microbes from next-generation sequencing data using breseq. Methods in Molecular Biology. 2014;1151:165–88.

43. Cairns BJ, Payne RJ. Bacteriophage therapy and the mutant selection window. Antimicrobial Agents and Chemotherapy. 2008;52(12):4344–50.

44. Abedon ST, Danis-Wlodarczyk KM, Wozniak DJ. Phage cocktail development for bacteriophage therapy: toward improving spectrum of activity breadth and depth. Pharmaceuticals (Basel). 2021;14(10).

45. Dettman JR, Sztepanacz JL, Kassen R. The properties of spontaneous mutations in the opportunistic pathogen Pseudomonas aeruginosa. BMC Genomics. 2016;17:27.

46. Heldal M, Bratbak G. Production and decay of viruses in aquatic environments. Marine Ecology Progress Series. 1991;72(3):205–12.

47. Kortright KE, Chan BK, Koff JL, Turner PE. Phage therapy: a renewed approach to combat antibiotic-resistant bacteria. Cell Host Microbe. 2019;25(2):219–32.

48. Weinbauer MG. Ecology of prokaryotic viruses. FEMS Microbiology Reviews. 2004;28(2):127–81.

49. Rampersad S, Tennant P. Chapter 3 - replication and expression strategies of viruses. In: Tennant P, Fermin G, Foster JE, editors. Viruses: Academic Press; 2018. p. 55–82.

50. Kleinman A. The mathematics of random mutation and natural selection for multiple simultaneous selection pressures and the evolution of antimicrobial drug resistance. Statistics in Medicine. 2016;35(29):5391–400.

51. Egido JE, Costa AR, Aparicio-Maldonado C, Haas PJ, Brouns SJJ. Mechanisms and clinical importance of bacteriophage resistance. FEMS Microbiology Reviews. 2022;46(1).

52. Mangalea MR, Duerkop BA. Fitness trade-offs resulting from bacteriophage resistance potentiate synergistic antibacterial strategies. Infection and Immunity. 2020;88(7).

53. Markwitz P, Olszak T, Gula G, Kowalska M, Arabski M, Drulis-Kawa Z. Emerging phage resistance in *Pseudomonas aeruginosa* PAO1 is accompanied by an enhanced heterogeneity and reduced virulence. Viruses. 2021;13(7).

54. Tiwari BR, Kim S, Rahman M, Kim J. Antibacterial efficacy of lytic *Pseudomonas* bacteriophage in normal and neutropenic mice models. Journal of Microbiology. 2011;49(6):994–9.

55. Leung CYJ, Weitz JS. Modeling the synergistic elimination of bacteria by phage and the innate immune system. Journal of Theoretical Biology. 2017;429:241–52.

56. Weld RJ, Butts C, Heinemann JA. Models of phage growth and their applicability to phage therapy. Journal of Theoretical Biology. 2004;227(1):1–11.

57. Dedrick RM, Freeman KG, Nguyen JA, Bahadirli-Talbott A, Smith BE, Wu AE, et al. Potent antibody-mediated neutralization limits bacteriophage treatment of a pulmonary Mycobacterium abscessus infection. Nature Medicine. 2021;27(8):1357–61.

58. Brun R, Reichert P, Künsch HR. Practical identifiability analysis of large environmental simulation models. Water Resources Research. 2001;37(4):1015–30.

